# Functional interplay between Mediator and RSC chromatin remodeling complex controls nucleosome-depleted region maintenance at promoters

**DOI:** 10.1101/2022.09.22.509083

**Authors:** Kévin M. André, Nathalie Giordanengo Aiach, Veronica Martinez-Fernandez, Leo Zeitler, Arach Goldar, Michel Werner, Cyril Denby Wilkes, Julie Soutourina

## Abstract

Chromatin organization is crucial for the transcriptional regulation in eukaryotes. Mediator is an essential and conserved coactivator thought to act in concert with chromatin regulators. However, it remains largely unknown how their functions are coordinated. Here, we provide evidence in the yeast *Saccharomyces cerevisiae* that Mediator establishes physical contact with RSC (Remodels the Structure of Chromatin), a conserved and essential chromatin remodeling complex that is crucial for nucleosome-depleted region (NDR) formation. We determined the role of Mediator-RSC interaction in their recruitment, nucleosome occupancy and transcription on a genomic scale. Mediator and RSC co-localize on wide NDRs of promoter regions, and specific Mediator mutations affect nucleosome eviction and TSS-associated +1 nucleosome stability. This work shows that Mediator contributes to RSC remodeling function to shape NDRs and maintain chromatin organization on promoter regions. It will help in our understanding of transcriptional regulation in the chromatin context relevant for severe diseases.

## Introduction

Eukaryotic DNA is compactly structured through chromatin organization which governs and influences all DNA transactions. Regulation of transcription is a key phenomenon of gene expression whose alterations lead to severe human pathologies. Multisubunit coregulator complexes constitute a key regulatory layer operating between specific transcription factors and basal RNA polymerase (Pol) II transcription machinery. These complexes can act on chromatin structure as chromatin modifiers or remodelers, or stimulate the assembly of the preinitiation complex (PIC). Some of them could have both functions. Many coregulator complexes are thought to cooperate. However, their functional interplay remains poorly understood.

Mediator is one of the essential and conserved coactivator complexes. The discovery of the Mediator complex three decades ago has revolutionized the field of transcription regulation (Kornberg, 2005). This key transcriptional coregulator transmits the regulatory signals from transcription factors to Pol II machinery and stimulates the PIC assembly in conjunction with the general transcription factors (Soutourina, 2018). Mediator is a huge complex (1.5 MDa) conserved in all eukaryotes and composed of 25 subunits in yeast, and of up to 30 subunits in human. Med17 is one of the ten essential Mediator subunits in yeast and mutations in this subunit globally affect Pol II transcription (Eyboulet et al., 2013; Eyboulet et al., 2015; Georges et al., 2019; Soutourina et al., 2011). In humans, given a key role of the Mediator complex in transcriptional regulation, mutations that affect Mediator subunits lead to various pathologies. Mediator is thought to activate transcription in concert with other coactivators including chromatin remodelers that alter the chromatin structure. However, the molecular mechanisms involved remain unknown. RSC (Remodels the Structure of Chromatin) is a multisubunit complex of the conserved SWI/SNF family of ATP-dependent chromatin remodelers (Clapier et al., 2017). It is composed of 16 subunits, is essential for cell viability in yeast and 10-fold more abundant than SWI/SNF complex, another remodeler of the same family. Two distinct human remodeling complexes BAF (Brg1 Associated Factors) and PBAF (Polybromo-associated BAF) are homologous to yeast SWI/SNF and RSC, respectively. A third complex closely related to BAF and PBAF, named ncBAF (non-canonical BAF complex) was recently identified in mammalian cells (Michel et al., 2018). An important role of BAF and PBAF complexes is highlighted by the fact that mutations of BAF/PBAF subunits have been linked to several diseases including cancers. ATP-dependent chromatin remodeling complexes and RSC in particular play a key role in the establishment of the promoter nucleosome-depleted regions (NDR) and nucleosome positioning on a genomic scale (Hartley and Madhani, 2009; Krietenstein et al., 2016; Kubik et al., 2018; Lorch et al., 2018; Yen et al., 2012; Zhang et al., 2011). Chromatin remodeling activity of RSC allows nucleosome sliding, destabilization or eviction (Clapier et al., 2016; Lorch et al., 1999; 2001). Furthermore, the Rsc3 and Rsc30 subunits of RSC belong to the Gal4-class of transcription factor-like proteins that recognize DNA motif preferentially found in promoter NDRs (Badis et al., 2008). Recent cryo-electron microscopy structural models of RSC bound to nucleosome revealed the modular organization of the RSC complex, structural details for its DNA and nucleosome recognition, as well as its role in NDR formation (Patel et al., 2019; Wagner et al., 2020; Ye et al., 2019). Sth1 is an essential catalytic subunit of RSC complex acting as ATP-dependent DNA translocase. Rsc8 subunit, essential for yeast viability, is present as a dimer and serves as a scaffold for the core structure of the RSC complex (Patel *et al*., 2019). Importantly, a cooperation between human Mediator, TFIID and specifically PBAF complex—human homolog of RSC—was reported in reconstituted *in vitro* transcription system with chromatin template (Lemon et al., 2001). While the molecular mechanism of chromatin remodeling by RSC complex is well established, the functional cooperation between RSC and Mediator, to our knowledge, has not been investigated. Understanding of the mechanisms involved in this cooperation represents an important and timely question in chromatin-related fields relevant for severe human diseases.

In this work, we investigate the RSC/Mediator functional interplay *in vivo* on a genomic scale. We have found that Mediator physically interacts with RSC chromatin remodeling complex and colocalizes with this remodeler on wide NDRs within promoter/regulatory regions. We identified Mediator mutations that affect RSC/Mediator interaction and RSC binding to the chromatin, and are lethal in combination with RSC mutations. Moreover, these Mediator mutations impair nucleosome distribution on RSC/Mediator-associated NDRs leading to the presence of an additional nucleosome on these regions. We propose that Mediator contributes to RSC remodeling function for nucleosome depletion to form and maintain NDRs on promoter regions. This work thus reveals that Mediator and RSC cooperate through their physical interaction for nucleosomal organization and transcription.

## Results

### Physical interaction between Mediator and RSC complexes

We previously performed a two-hybrid screening for Mediator subunits with a yeast genomic library that allowed to characterize protein-protein interactions within the complex (Guglielmi et al., 2004). Additional interactions between Mediator subunits and other nuclear proteins were revealed and reported for Rad3 and Rad2 (Esnault et al., 2008; Eyboulet *et al*., 2013). They also include interactions with RSC subunits that were not published. In particular, the Med17 Mediator head module subunit fused to the Gal4 DNA binding domain (G_BD_) interacts in the two-hybrid assay with a Rsc8 RSC subunit fragment (amino acids 248-465) fused to the Gal4 activation domain (G_AD_) (**Figure 1A**). We also showed that this interaction is maintained with the full-length Rsc8, as well as when Rsc8 was fused to G_BD_ and Med17 was fused to G_AD_. In addition to the contact between Med17 and Rsc8, Med21 Mediator subunit fused to the G_BD_ interacts with Rsc3 fragment (amino acids 588-695) and weakly with Rsc30 fragment (amino acids 644-883) (**Supplementary Figure S1**). Further study was focused on the Med17-Rsc8 interaction. To identify the Med17 domain that is required for its interaction with Rsc8, we analyzed different N- and C-terminal Med17 truncation mutants for their interaction with Rsc8 full-length and Rsc8 fragment (**Figure 1B**). We conclude that Med17 domain (amino acids 346–687) is necessary and sufficient for Med17 interaction with Rsc8.

**Figure 1.**
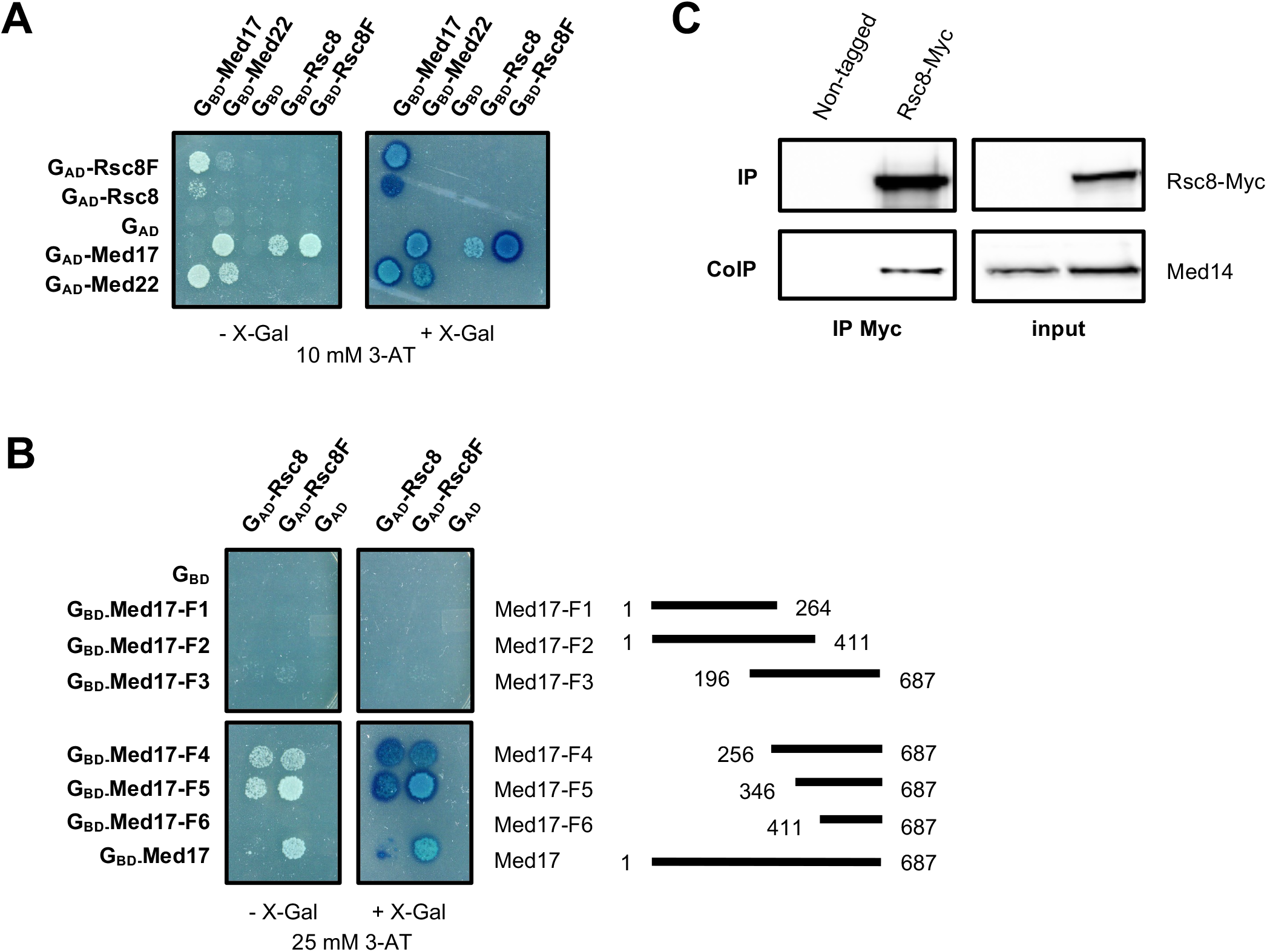
Mediator and RSC physically interact. (**A**) Gal4 DNA-binding domain (G_BD_), alone or in fusion with full length Med17, Med22, Rsc8 or Rsc8 fragment (Rsc8F: amino acids 248-465), was co-expressed with Gal4 activation domain (G_AD_) alone or in fusion with Rsc8, Rsc8 fragment, Med17 or Med22. Interaction between Med17 and Med22 subunits of Mediator was included as a positive control. Yeast cells were spotted on SD+2A medium supplemented with 10 mM 3-AT, grown for 3 days (left panel) and then stained with X-Gal for 24h (right panel). (**B**) Gal4 DNA-binding domain (G_BD_), alone or in fusion with either full length Med17 or truncations of Med17 (termed Med17 fragments (F) 1 to 6), was co-expressed with Gal4 activation domain (G_AD_) alone or in fusion with full-length Rsc8 or Rsc8 fragment (Rsc8F). Yeast cells were spotted on SD+2A medium supplemented with 25 mM 3-AT, grown for 3 days (left panel) and then stained with X-Gal for 24h (right panel). The boundaries of the Med17 fragments in amino acids are represented on the right. (**C**) Western blot analysis of Mediator-RSC interaction. Crude extracts were prepared from yeast WT strain expressing a tagged version of Rsc8 (Myc) following a 45-min shift at 37°C, after reaching exponential phase at 30°C. A strain carrying non-tagged Rsc8 was used as a negative control. Samples were immunoprecipitated with anti-Myc antibodies. Immunoprecipitated proteins and inputs were analyzed by western blotting with anti-Myc and anti-Med14 antibodies (n=3). See also Figure S1.

The contact between Mediator and RSC complexes was confirmed by coimmunoprecipitation (co-IP) experiments with crude extracts of a yeast strain expressing Rsc8-Myc from its native promoter (**Figure 1C**). Our results show that Mediator coimmunoprecipitates with Rsc8 in crude extracts. Indeed, the Med14 subunit of Mediator was detected by Western blotting when Rsc8-Myc was used to immunoprecipitate RSC complex.

### Mediator and RSC colocalize on the promoter regions

To compare how Mediator and RSC are distributed on the yeast genome, we performed ChIP-seq experiments for Mediator (Med14 subunit), as well as RSC (Rsc8 and Sth1 subunits). Input DNA and DNA from ChIP with a non-tagged strain were used as negative controls. Yeast strains carrying C-terminal Myc-tagged version of Med14, Rsc8 and Sth1 were constructed and utilized in ChIP-seq experiments. In accordance with previous studies, heat map of tag density centered on transcription start sites (TSS) shows Mediator and RSC enrichment on promoter regions (**Figure 2A**). Specific examples of ChIP-seq profiles illustrate Mediator and RSC enrichment on promoter regions of Pol II-transcribed genes (**Supplementary Figure S2A**). Analysis of Mediator (Med14 subunit) and RSC (Rsc8 and Sth1 subunits) enrichment peaks on intergenic regions corresponding to upstream activating sequences (UAS) revealed that most of the Med14 peaks (242 out of 313) overlap with RSC peaks (**Figure 2B** and **Supplementary Figure S2B**). Heat maps of tag density generated for Mediator and RSC centered on Med14, Rsc8 and Sth1 enrichment peaks clearly illustrate Mediator colocalization with RSC (**Figure 2C** and **Supplementary Figure S2C**). Analysis on intergenic regions showed a strong correlation between Rsc8 and Sth1 sets of data (Spearman’s correlation coefficient (SCC) equal to 0.89), as expected for the subunits of the same remodeling complex, and moderate correlation between Rsc8/Sth1 and Mediator occupancies (SCC equal to 0.54 and 0.61, respectively) (**Supplementary Figure S2D**).

**Figure 2.**
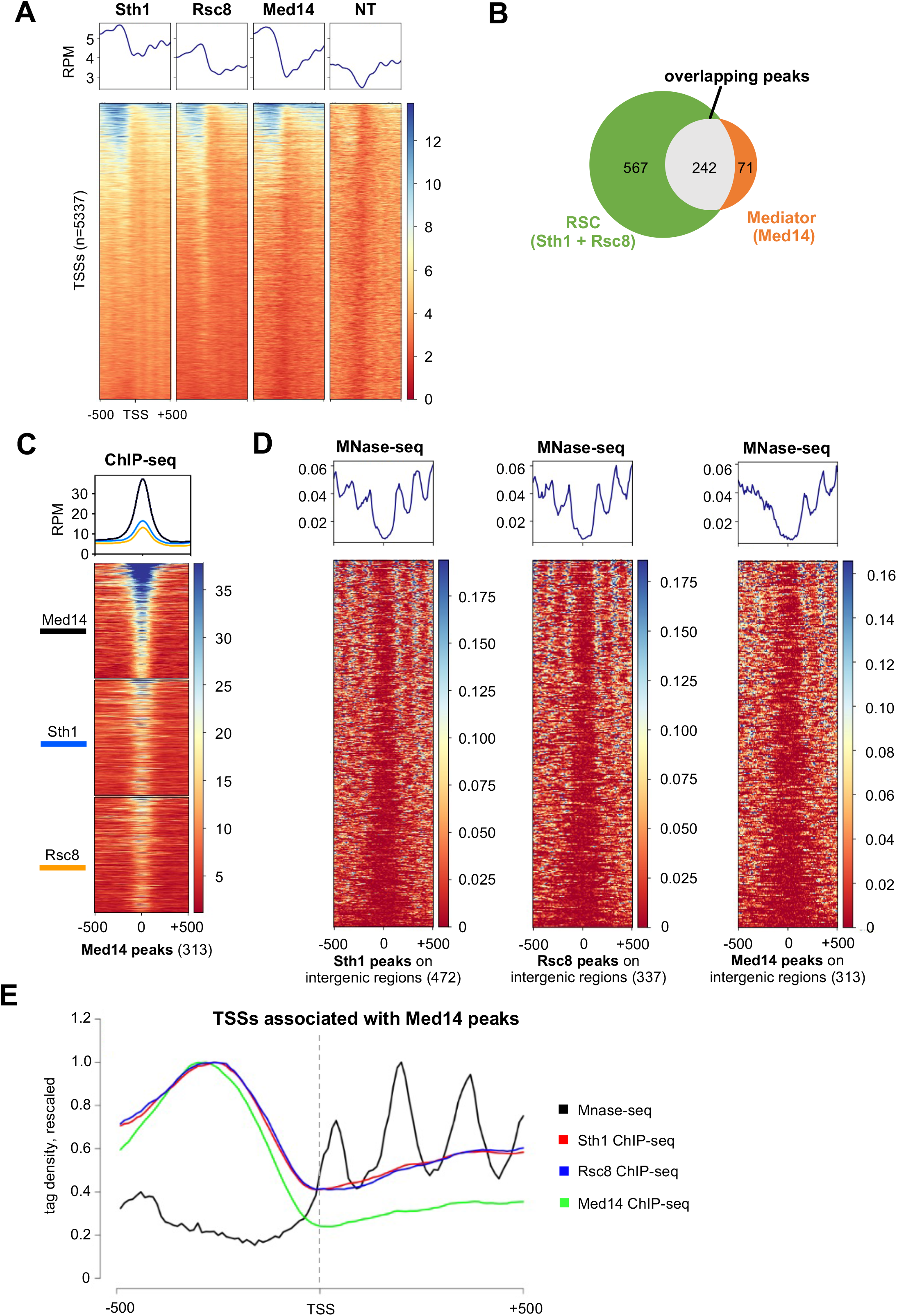
Mediator and RSC colocalize on NDRs within promoter regions. (**A**) Heat maps and average profiles of Sth1-Myc, Rsc8-Myc and Med14-Myc ChIP-seq signals centered on TSSs (−500 bp to +500 bp). Negative control was obtained from a Myc IP in a non-tagged strain (NT). Signals are expressed as the normalized read counts. (**B**) Venn diagram illustrating the number of Mediator (Med14) peaks that overlap or not with RSC peaks, which are defined as overlapping peaks between Sth1 and Rsc8 plus the unique ones for each protein. (**C**) Heat maps and average profiles of Med14-Myc, Sth1-Myc and Rsc8-Myc centered on Med14 peaks within intergenic regions (−500 bp to +500 bp). Signals are expressed as the normalized read counts. (**D**) Heatmaps and average profiles of MNase-digested DNA centered on Sth1, Rsc8 or Med17 peaks within intergenic regions (−500 bp to +500 bp). The number of peaks for each protein is indicated in brackets. Signals are expressed as the normalized read counts. (**E**) Sth1, Rsc8, Med14 and MNase profiles centered on TSSs associated to Med14 peaks (−500 bp to +500 bp). The four profiles were normalized for comparison by rescaling their maximum values to 1. See also Figure S2.

To further analyze RSC and Mediator enrichment peaks on intergenic regions, we compared their location with the nucleosome occupancy determined by MNase-seq. Heat maps of tag density generated by MNase-seq were centered on Med14, Rsc8 and Sth1 enrichment peaks, showing colocalization of RSC and Mediator peaks with nucleosome-depleted regions (NDR) (**Figure 2D**). The center of NDRs with minimum of nucleosome occupancy colocalizes with Med14, Rsc8 and Sth1 enrichment peaks. By comparing Mediator and RSC ChIP-seq profiles together with nucleosome distribution determined by MNase-seq in relation to TSS, we showed that Mediator and RSC are enriched on NDRs upstream of TSSs (**Figure 2E** and **Supplementary Figure S2E**). Taken together, our results show that Mediator and RSC colocalize on promoter regions of Pol II-transcribed genes that correspond to NDR.

### RSC mutations affect nucleosome distribution and Mediator occupancy genome-wide

Since Mediator and RSC physically interact and share their binding location on promoter regions, we decided to take advantage of specific mutants in these two complexes to investigate their interrelationships for chromatin binding, nucleosomal organization and transcription. For RSC, we focused on the temperature-sensitive mutants in Rsc8 subunit that interacts with Mediator and Sth1 catalytic subunit of the complex. We analyzed the effect of *rsc8-ts* (G326D, I417F, K446M) and *sth1-ts* (P646L, F655L, D657G, N1077S, F1125L) mutations (Li et al., 2011) compared to the wild type (WT) strains after a shift to non-permissive temperature. Genome-wide nucleosome distribution was analyzed by MNase-seq and Pol II, Mediator (Med14-Myc tagged subunit) and RSC (Rsc8 or Sth1-Myc tagged subunits) distribution were determined by ChIP-seq. Consistent with an essential role of this chromatin remodeling complex in nucleosome positioning and NDR formation on promoters and previously reported MNase-seq profiles in RSC-depleted cells (Kubik *et al*., 2018), *rsc8* and *sth1* mutations led to global changes in nucleosome distribution with an upstream shift of nucleosomes on transcribed regions and a downstream shift of those on promoter regions (**Figure 3A**). This results in a significant decrease in the NDR length on promoter regions (**Figure 3B**). Focusing on RSC Sth1 enrichment peaks associated to NDR regions, heat maps of MNase-seq data illustrate major changes in *rsc8-ts* mutant compared to the WT strain (**Figure 3C**). In both mutants, a global effect on Pol II transcription was observed, as illustrated by a 3-fold decrease in Pol II ChIP-seq density on transcribed regions (**Supplementary Figure S3A**). Heat maps centered on Mediator or RSC peaks within intergenic regions as defined in the WT context show that *rsc8-ts* mutations affect both Mediator and RSC occupancy, leading to a decrease in the presence of Med14 and Sth1 on chromatin (**Figure 3D** and **Supplementary Figure S3B**). Our analysis of *sth1-ts* mutant reveals, on the contrary, a stabilization of Mediator on enrichment peaks of both complexes in this mutant compared to the WT and no major effect on Rsc8 occupancy (**Figure 3D** and **Supplementary Figure S3B**). ChIP followed by qPCR was also performed on selected regions (**Supplementary Figure S4**). Interestingly, these changes in Mediator occupancy in RSC mutants were accompanied by changes in physical interaction between Mediator and RSC. We observed a decrease in coimmunoprecipitation signal between Mediator and RSC complexes in *rsc8-ts* mutant and, on the contrary, an increase for *sth1-ts* mutant (**Figure 3E**). Taken together, our results show that RSC mutations affect global nucleosome organization and NDR formation on promoter regions. Moreover, we reveal their effect on Mediator occupancy associated with changes in physical interaction between RSC and Mediator.

**Figure 3.**
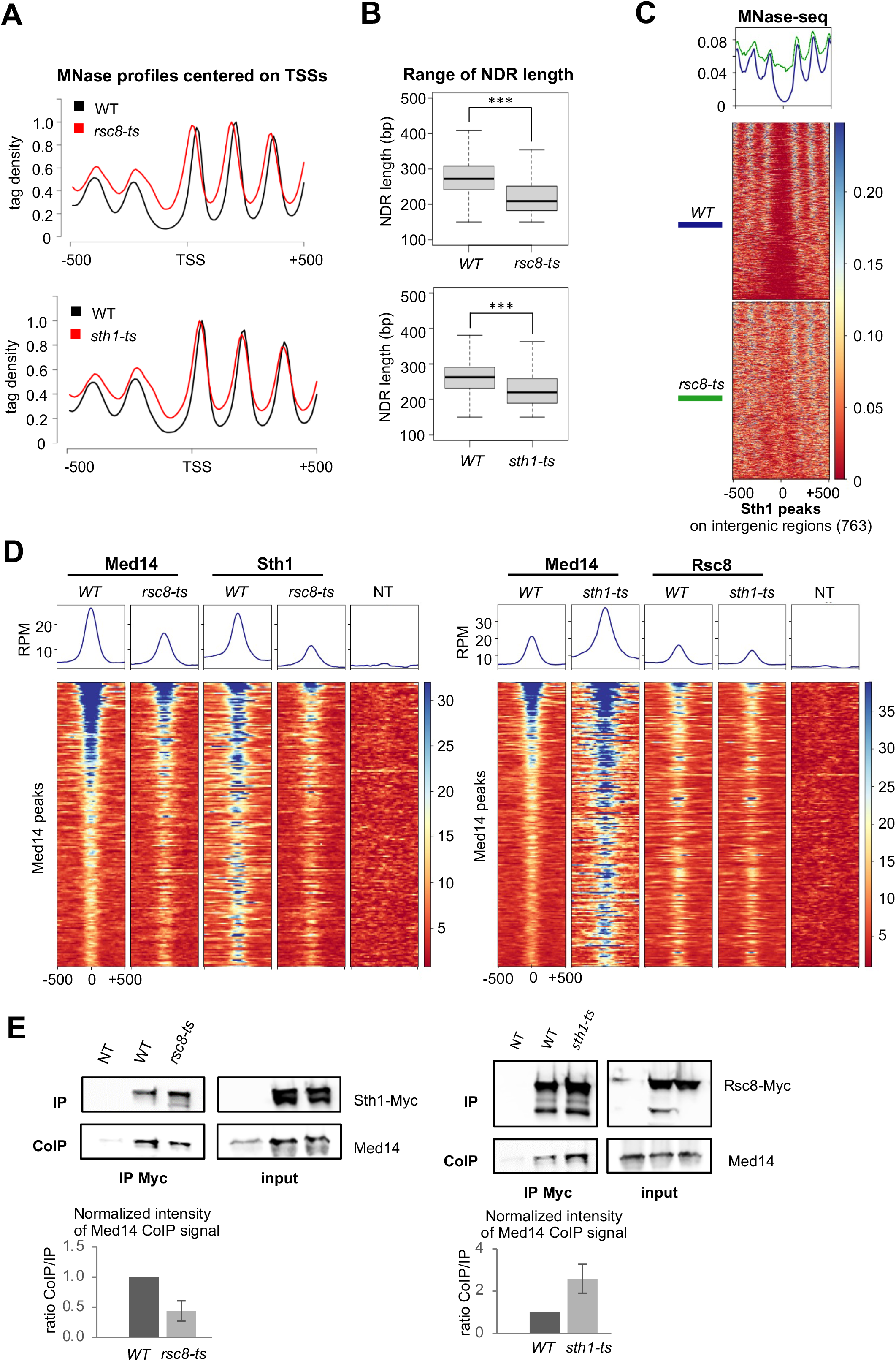
RSC mutations impair the NDR architecture and the Mediator-RSC interaction. (**A**) Average MNase profiles centered on TSSs (n=5337, −500 bp to +500 bp), in WT, *rsc8-ts* and *sth1-*ts strains. The profiles were normalized for comparison by rescaling their maximum values to 1. (**B**) Boxplot showing the range of NDR lengths in WT, *rsc8-ts* and *sth1-ts* strains. The three asterisks represent a significant difference between the WT and the mutant at P-value < 0.001 in a Wilcoxon test. (**C**) Heat maps and average MNase profiles centered on Sth1 peaks within intergenic regions (n=763, −500 bp to +500 bp). Signals are expressed as the normalized read counts. (**D**) Left panel: Heat maps and average profiles of Med14-Myc and Sth1-Myc ChIP-seq signals in WT or *rsc8-ts* centered on Med14 WT peaks (n=187, −500 bp to +500 bp). Right panel: Heat maps and average profiles of Med14-Myc and Rsc8-Myc ChIP-seq signals in WT or *sth1-ts* centered on Med14 WT peaks (n=209, −500 bp to +500 bp). Negative control was obtained from a Myc IP in a non-tagged strain (NT). Signals are expressed as the normalized read counts. (**E**) Western blot analysis of Mediator-RSC interaction. Crude extracts were prepared from yeasts expressing tagged versions of Sth1 or Rsc8 (Myc) in the WT or mutant strains (*rsc8-ts* in the left panel and *sth1-ts* in the right panel, respectively) following a 6-hour or 2.5-hour shift at 37°C (for *rsc8-ts* and *sth1-ts*, respectively), after reaching exponential phase at 30°C. Samples were immunoprecipitated with anti-Myc antibodies. Immunoprecipitated proteins and inputs were analyzed by western blotting with anti-Myc and anti-Med14 antibodies. The intensity of immune staining for the coimmunoprecipitated Med14 signals was normalized against the immunoprecipitation and input signals (bottom panels). The mean values and standard deviations (indicated by error bars) of at least three independent experiments are shown. The asterisk represents a significant difference between the WT and the mutant at P-value <0.05 in a Student t-test. See also Figures S3, S4.

### Specific Mediator mutations affect NDR formation/maintenance

To further investigate relationships between Mediator and RSC, we focused on well-studied temperature-sensitive mutants in Med17 Mediator subunit (Eyboulet *et al*., 2013; Eyboulet *et al*., 2015; Georges *et al*., 2019) that interacts with RSC. We combined *med17* mutations with *rsc8-ts* or *sth1-ts* RSC mutations and showed that *med17-140* (Q444P/M442L) was colethal in both RSC mutant contexts (**Figure 4A**). *med17-444* (Q444P) mutation did not lead to any synthetic phenotype in combination with *rsc8-ts* and showed a slower growth in *sth1-ts* context. Allele-specific lethality between Mediator and RSC mutations is consistent with the physiological importance of their interaction. Further analysis of physical interaction between Mediator and RSC, as well as genome-wide distribution of nucleosomes, Pol II, Mediator and RSC was focused on *med17-140* mutant compared to the WT strain. *med17-444* mutant was also included in the analysis for comparison. As previously described, both *med17-444* and *med17-140* mutations led to a strong transcriptional defect with a global decrease in Pol II occupancy compared to the WT (**Supplementary Figure S5A**).

**Figure 4.**
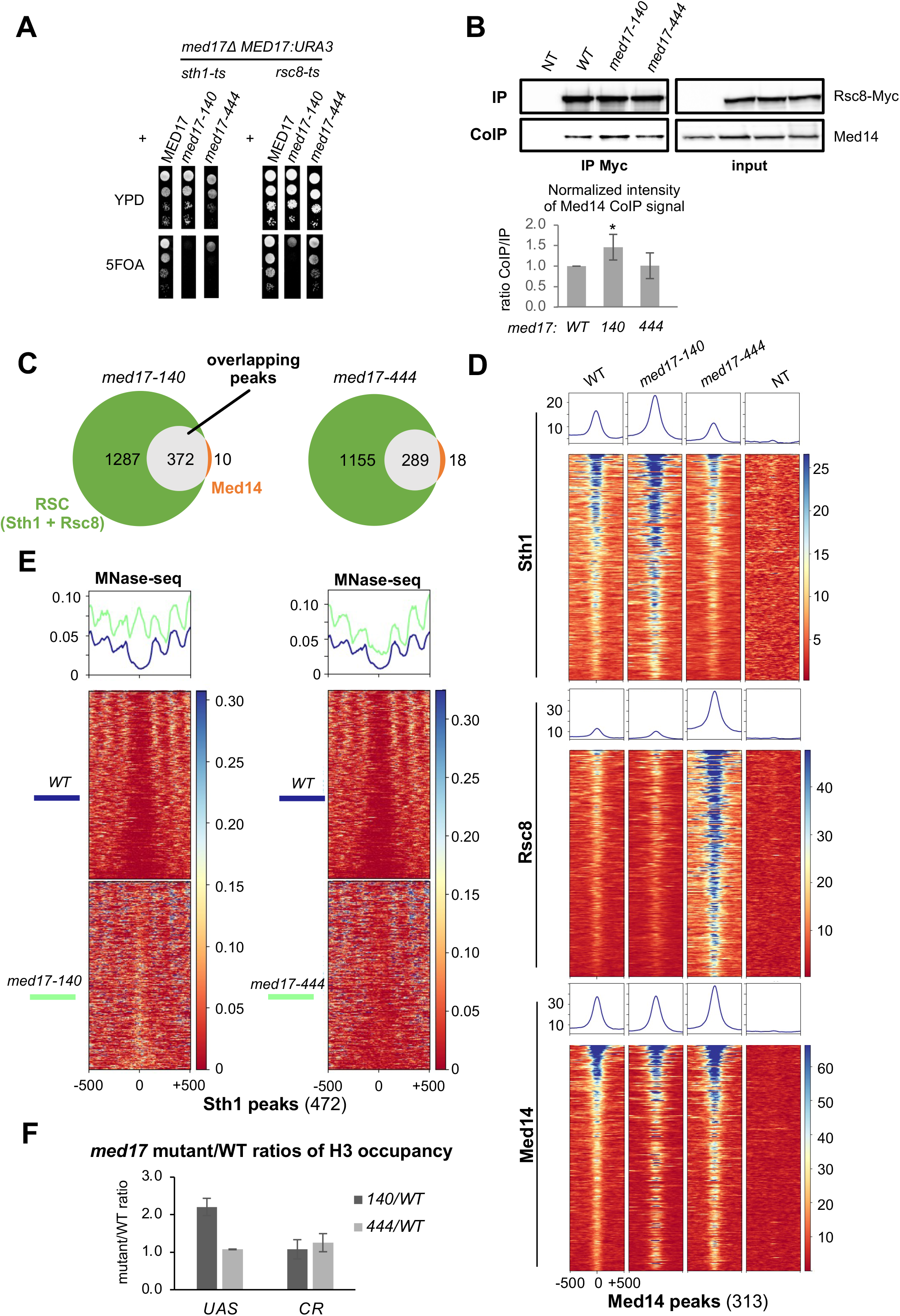
Mediator-RSC interaction and NDRs are disturbed in *med17-140*. (**A**) Synthetic phenotypes between *rsc* and *med17* mutations. RSC mutants (*sth1-ts* and *rsc8-ts*) have the deletion of *MED17* gene, complemented with a plasmid containing WT *MED17:URA3. rsc* mutants were transformed by plasmids containing either WT or mutant (*med17-140, med17-444*) versions of Med17. The MED17 WT copy (*MED17:URA3*) was counterselected by spotting the strains on 5-FOA medium. (**B**) Western blot analysis of Mediator-RSC interaction. Crude extracts were prepared from yeast cells expressing tagged versions of Rsc8 (Myc) in the WT or mutant strains (*med17-140* and *med17-444*) following a 45-min shift at 37°C, after reaching exponential phase at 30°C. Samples were immunoprecipitated with anti-Myc antibodies. Immunoprecipitated proteins and inputs were analyzed by western blotting with anti-Myc and anti-Med14 antibodies. The intensity of immune staining for the coimmunoprecipitated Med14 signals was normalized against the immunoprecipitation and input signals. The mean values and standard deviations (indicated by error bars) of three independent experiments are shown. The asterisk represents a significant difference between the WT and the mutant at P-value <0.05 in a Student t-test. (**C**) Venn diagrams illustrating the number of Med14 peaks that overlap or not with RSC peaks in *med17-140* (left panel) and *med17-444* (right panel). RSC peaks are defined as Sth1 and Rsc8 overlapping peaks plus the ones that are specific for each protein. (**D**) Heat maps and average profiles of Sth1 (top), Rsc8 (middle) and Med14 (bottom) ChIP-seq signals centered on Med14 WT peaks within intergenic regions (n=313, −500 bp to +500 bp). Negative control was obtained from a Myc IP in a non-tagged strain (NT). Signals are expressed as the normalized read counts. (**E**) Heat maps and average MNase profiles centered on Sth1 peaks within intergenic regions (n=472, −500 bp to +500 bp) in the WT, *med17-140* (left panel) and *med17-444* (right panel) strains. Signals are expressed as the normalized read counts. (**F**) Ratios of mutant/WT H3 occupancy for *med17- 140* (dark gray) and *med17-444* (light gray) on both upstream activating sequences (UAS) and coding regions (CR). Average ratios were calculated from quantitative PCR data (see **Supplementary Figure S7**) observed on three UASs and three CRs. See also Figures S5-7.

We examined the effect of *med17* mutations on Mediator interaction with RSC by CoIP experiments. We found a slight but significant increase (P-value <0.05 in a Student t-test) in RSC/Mediator CoIP in crude extract from *med17-140* mutant compared to the WT and *med17-444* mutant (**Figure 4B**). This observation suggests that *med17-140* mutation modifies the physical interaction between Mediator and RSC.

We then investigated by ChIP-seq whether Mediator and RSC occupancies were affected in *med17* mutants. Our analysis of Mediator (Med14 subunit) and RSC (Rsc8 and Sth1 subunits) enrichment peaks on intergenic regions showed that the majority of the Med14 peaks overlap with RSC peaks in *med17-140* and *med17-444* mutants (372 out of 382 or 289 out of 307, respectively) (**Figure 4C** and **Supplementary Figure S5B**). Similar to the WT strain, Mediator and RSC colocalization was clearly observed in *med17* mutants (**Supplementary Figure S5C**). A strong correlation between Rsc8 and Sth1 data sets was also maintained in both mutants (SCC equal to 0.93 and 0.90) (**Supplementary Figure S5D**). Correlation between Rsc8/Sth1 and Mediator occupancies observed in the WT was reduced in *med17-444* (SCC equal to 0.46 and 0.54, respectively) and even more in *med17-140* (SCC equal to 0.40 and 0.50, respectively) (**Supplementary Figure S5D**).

Our analysis revealed that *med17* mutations affect RSC occupancy. Heat maps centered on Mediator peaks on intergenic regions as defined in the WT context show that *med17-140* mutations lead to increased presence of Sth1 on the Mediator peaks (**Figure 4D** and **Supplementary Figure S5E**). In *med17-444* mutant, Rsc8 occupancy was increased (**Figure 4D** and **Supplementary Figure S5E**). For Mediator occupancy, no changes in distribution were observed in *med17-140* and only a slight increase was seen in *med17-444* mutants compared to the WT. ChIP followed by qPCR was also performed on selected regions for core promoters, UASs and gene bodies and confirmed our genome-wide analysis, illustrating effects of *med17* mutations on Pol II, Mediator and RSC occupancies (**Supplementary Figure S6**).

We then analyzed the nucleosome occupancy in *med17* mutants and observed a great effect of *med17-140* mutations on MNase-seq signal on promoter regions (**Figure 4E** and **Supplementary Figure S5F**). Metagene analysis and heat maps centered on RSC and Mediator enrichment peaks clearly show that NDR regions associated to these peaks were dramatically changed in *med17-140* compared to the WT and *med17-444* mutant. An additional peak was observed in *med17-140* mutant at the center of NDRs. To confirm that this increase in MNase-seq signal corresponds to nucleosome occupancy, we performed histone H3 ChIP analysis. We showed that H3 occupancy of UASs was increased by 2-fold in *med17-140* mutant compared to the WT and no changes were observed in *med17-444* mutant (**Figure 4F** and **Supplementary Figure S7A**). This effect was specific to UASs and was not observed for other tested regions including core promoters and gene bodies. Examples of MNase-seq profiles around selected UASs clearly illustrate a specific increase in nucleosome occupancy of these regions in *med17-140* mutant compared to the WT and *med17-444* mutant (**Supplementary Figure S7B**). Taken together, our results suggest that Mediator is involved in the maintenance of NDRs associated to RSC and itself.

### Specific Mediator mutations affect the maintenance of wide NDRs bound by Mediator and RSC

To further investigate the Mediator effect on NDR maintenance, we classified Mediator and RSC enrichment peaks in two categories: peaks shared between the two complexes and unique peaks enriched only by RSC. Metagene analysis and heat maps in *med17-140* show the most important increase in nucleosome occupancy on NDRs associated to shared Mediator/RSC peaks compared to unique peaks (**Figure 5A** and **Supplementary Figure S8A**). We then performed metagene analysis of rescaled tag density centered on TSS that revealed the most pronounced changes in MNase-seq profiles on NDR upstream of TSS associated with shared Mediator/RSC peaks (**Figure 5B** and **Supplementary Figure S8B**). In addition, the MNase-seq signal at +1 nucleosome position was reduced in *med17-140* mutant in comparison with +2 and +3 nucleosomes, suggesting that Mediator mutations lead to filling of NDR and destabilization of the TSS-associated +1 nucleosome. We also noticed that shared Mediator/RSC peaks are characterized by a higher RSC occupancy compared to the unique RSC peaks and a higher Sth1 stabilization in *med17-140* mutant with respect to the WT (**Figure 5C, D** and **Supplementary Figure S8C, D**). Rsc8 occupancy decreased in *med17-140* mutant compared to the WT, while Mediator occupancy remained close to the WT level (**Supplementary Figure S8C, D**).

**Figure 5.**
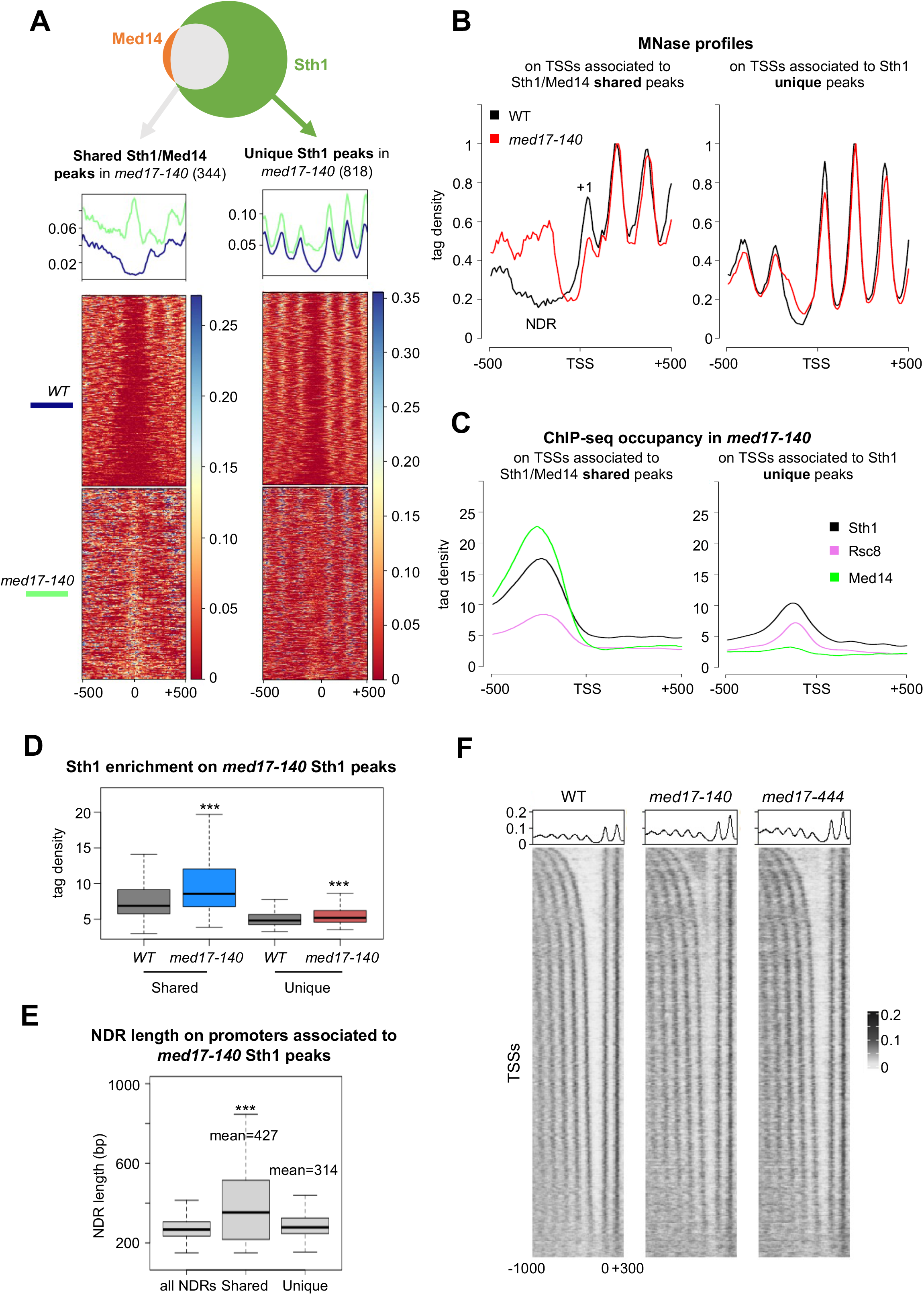
NDRs are disturbed in *med17-140* on promoter regions enriched by Mediator and RSC. (**A**) Heat maps and average MNase profiles centered on *med17-140* Sth1 peaks that overlap (left panel) or not (right panel) with Med14 peaks (n=344 and 818, respectively, −500 bp to +500 bp) within intergenic regions in the WT and *med17-140* strains. Signals are expressed as the normalized read counts. (**B**) Average MNase profiles centered on TSSs associated to Sth1/Med14-shared peaks (n=344) or unique Sth1 peaks (n=818), from −500 bp to +500 bp, in WT and *med17-140*. The profiles were normalized for comparison by rescaling their maximum values to 1. NDR: nucleosome-depleted region. +1: nucleosome +1. (**C**) Sth1, Rsc8 and Med14 profiles centered on TSSs associated to shared Med14/Sth1peaks (left panel, n=344) or unique Sth1 peaks (right panel, n=818), −500 bp to +500 bp. (**D**) Boxplots showing the Sth1 enrichments in WT and *med17-140* on *med17-140* Sth1 peaks that overlap (termed shared) or not (termed unique) with Med14 peaks. The three asterisks represent a significant difference between the WT and the mutant at P-value < 0.001 in a Wilcoxon test. (**E**) Boxplots showing the NDR lengths on promoters associated to both Sth1 and Med14 (middle) or Sth1 only (right). The three asterisks represent a significant difference between the Sth1/Med14-associated NDRs and all NDRs or Sth1-associated NDRs at P-value < 0.001 in a Wilcoxon test. (**F**) Heat maps and average MNase profiles centered on TSSs (n=5337, −1000 bp to +300 bp) in the WT, *med17-140* and *med17-444* strains. TSS regions were sorted according to the size of the associated NDR. Signals are expressed as the normalized read counts. See also Figures S8, S9.

We then compared the length of NDRs associated with shared Mediator/RSC peaks and unique RSC peaks (**Figure 5E** and **Supplementary Figure S8E, F**). Our analysis showed that shared Mediator/RSC peaks are associated with wider NDRs (mean size equal to 427 bp in comparison with 314 for unique peaks) and larger range of NDR sizes (**Figure 5E** and **Supplementary Figure S8E, F**). When nucleosome occupancy data for the WT and *med17* mutants were visualized by ordering promoter-associated NDRs by their length for all Pol II-transcribed genes, a clear increase in nucleosome occupancy was observed in *med17-140* for wider NDRs as defined in the WT (**Figure 5F**).

As described above, Pol II transcription was globally affected in *med17-140* mutant (**Supplementary Figure S5A**). Our Pol II ChIP-seq analysis showed that genes associated with shared RSC/Mediator peaks correspond to higher transcription level compared to genes associated with unique RSC peaks (**Supplementary Figure S9**). A decrease in transcriptional level in *med17-140* mutant was comparable for both groups of genes.

Taken together, our results demonstrate that Mediator mutations affecting RSC/Mediator interaction and RSC binding to the chromatin, lead to a substantial change in nucleosome occupancy on wide NDRs associated with RSC and Mediator. This suggests that Mediator is involved in RSC-dependent NDR formation or maintenance on promoter/regulatory regions.

## Discussion

In this study, we addressed the key question on the functional interplay between two essential coregulator complexes acting in transcription and chromatin organization, Mediator and RSC chromatin remodeler. **Figure 6** illustrates, based on our findings, how interrelationships between these two complexes contribute to nucleosome organization on promoter regions through NDR formation or maintenance. Our four main outcomes can be summarized as follows. (i) We show that Mediator and RSC physically interact and identify specific subunits of these complexes, in particular Med17 and Rsc8, that are in contact. (ii) Furthermore, Mediator and RSC co-localize on intergenic regions, with RSC being enriched on Mediator peaks and vice-versa. We then show that Mediator and RSC are enriched on NDRs upstream the TSS and that shared RSC/Mediator peaks are associated with wide NDRs. (iii) Taking advantage of yeast genetics, our study shows that mutations in RSC complex affect chromatin occupancy of Mediator and vice-versa. Consistent with a functional link between the two coregulators, specific Mediator and RSC mutations are co-lethal. (iv) Mediator mutations (*med17-140*) that affect RSC/Mediator interaction and RSC binding to the chromatin lead to a great change in nucleosome occupancy on RSC/Mediator-associated NDRs and the presence of an additional nucleosome on these regions. Moreover, the +1 nucleosome occupancy was reduced in this mutant, suggesting that Mediator mutations lead to destabilization of the TSS-associated +1 nucleosome. Taken together, our results reveal that Mediator and RSC cooperate through their physical interaction for nucleosomal organization and transcription. We propose that Mediator is involved in RSC-dependent NDR formation or/and maintenance on promoter/regulatory regions by contributing to RSC remodeling function for nucleosome eviction within NDRs and stable positioning of +1 nucleosome.

**Figure 6.**
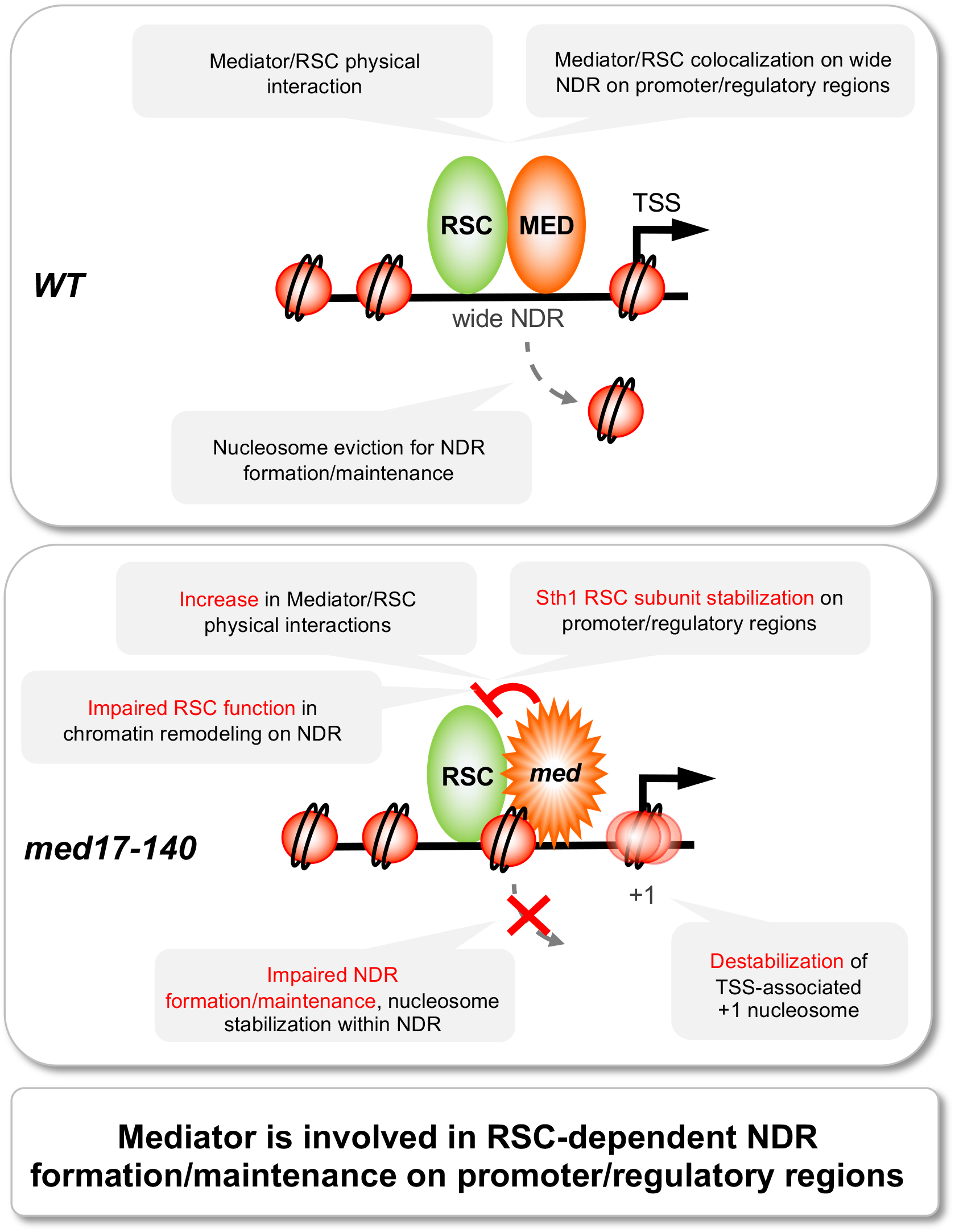
Mediator regulates RSC function in NDR formation/maintenance. In the WT context (upper panel), RSC and Mediator physically interact and colocalize on promoter/regulatory regions. Their cooperation contributes to RSC function in nucleosome eviction for wide NDR formation and stable positioning of TSS-associated +1 nucleosomes. In *med17-140* Mediator mutant (lower panel), physical interaction between RSC and Mediator is increased and Sth1 RSC subunit is stabilized on promoter/regulatory regions that disturb RSC remodeling activities. As a consequence, nucleosomes within NDRs are not evicted and +1 nucleosomes are destabilized. We propose that Mediator is involved in RSC-dependent NDR formation/maintenance on promoter/regulatory regions.

To our knowledge, this work provides the first evidence that Mediator contributes to nucleosome distribution and NDR formation or/and maintenance on promoter/regulatory regions in cooperation with RSC complex. The effect of Mediator mutations on nucleosome occupancy could not be explained by a global decrease in transcription *per se* since it was observed only for *med17-140* but not for *med17-444* with a similar decrease in Pol II occupancy. Moreover, it was previously shown that a decrease of transcription in *rpb1-1* Pol II mutant did not lead to any differences in nucleosome distribution (Hartley and Madhani, 2009). Mechanisms of chromatin remodeling activity of RSC to form wide NDRs were investigated in several studies (Brahma and Henikoff, 2019; Hartley and Madhani, 2009; Kubik et al., 2015; Kubik *et al*., 2018). Moreover, a recent thermodynamic model shows that RSC is the most significant contributor of NDR, allowing to predict the correct size for the large proportion of NDRs (Kharerin and Bai, 2021). Indeed, RSC was shown to be required for the normal positioning of NDR-flanking nucleosomes on the majority of promoter regions (Hartley and Madhani, 2009; Krietenstein *et al*., 2016; Kubik *et al*., 2015). An additional participation of Abf1 and Reb1 general regulatory factors (GRFs) on a subset of promoter regions was reported (Hartley and Madhani, 2009; Kubik *et al*., 2015), even though the exact relationships between RSC and GRF actions on the chromatin architecture of promoter regions remain unclear. To explain the properties of nucleosomes at promoter NDRs, it has been proposed that unstable or “fragile” nucleosomes are present at promoter regions of 40% of Pol II-transcribed genes corresponding to near 2000 genes (Kubik *et al*., 2015). More recently, RSC was shown to bind to nucleosomes at wide NDRs and proposed to mobilize and destabilize them (Brahma and Henikoff, 2019). Based on these results, fragile nucleosomes were suggested to be transient intermediates of RSC remodeling (Brahma and Henikoff, 2019). We propose that Mediator contributes to RSC-dependent remodeling mechanisms at promoter regions.

Our results suggest that an increase in physical interaction between Mediator and RSC, and a stabilization of Sth1 RSC subunit on wide NDRs can affect RSC function in nucleosome eviction on these regions, without major effect on positioning of nucleosomes +2 and +3. Interestingly, *in vitro* study revealed that an enhanced RSC-nucleosome interaction due to histone acetylation by SAGA has an inhibitory role on RSC function in nucleosome eviction, without affecting all RSC activities (Lorch *et al*., 2018). This means that an enhancement in RSC interactions with its partners such as nucleosomes can locally impact some of the RSC remodeling activities.

It was previously hypothesized that the function of RSC in nucleosome eviction at promoter regions can differ from its function in +1 and −1 nucleosome positioning or that RSC bound to a nucleosome can transit between +1 and −1 nucleosome positions, with RSC-associated particle at NDR being a transient intermediate (Brahma and Henikoff, 2019). Remarkably, recent studies using single-molecule imaging reveal that RSC, together with other chromatin remodelers, frequently undergoes short-lived chromatin interactions and suggest a dynamic competition between PIC assembly and the +1 nucleosome (Kim et al., 2021; Nguyen et al., 2021). In any case, it is strongly assumed that the NDR formation/maintenance and positioning of flanking +1 and −1 nucleosomes should be closely related. In this work we show that Mediator *med17-140* mutations lead to filling of wide RSC/Mediator-associated NDRs by precluding nucleosome eviction and to destabilization of the TSS-associated +1 nucleosome. It is well established that RSC together with other chromatin remodelers participates in positioning of +1 nucleosome (Krietenstein *et al*., 2016; Kubik *et al*., 2018). Interestingly, *in vitro* study demonstrated that enhanced RSC-histone interaction stabilizes +1 nucleosome by preventing its removal by RSC (Lorch *et al*., 2018). The +1 nucleosomes are characterized by high turnover (Dion et al., 2007); they are asymmetric, partially unwrapped and associated to RSC (Ramachandran et al., 2015). We propose that Mediator mutations can affect RSC activities leading to destabilization of +1 nucleosome through impaired nucleosome turnover or its less defined-positioning. Importantly, these nucleosomes have a key role for PIC assembly and transcription initiation since they are associated with TSS. PIC and nucleosomes can be assembled in competitive or cooperative manner depending on the promoters (Rhee and Pugh, 2012). It is well established that Mediator is essential in PIC assembly and we previously suggested that this role is related to chromatin architecture in terms of nucleosome occupancy and dynamics (Eychenne et al., 2016; Soutourina, 2018).

RSC is also known for its role in 3D chromatin organization. This complex is enriched on strong boundaries between chromosomal interacting domains (CIDs), and mutations in RSC subunits affect the folding of the yeast genome (Hsieh et al., 2015). Mediator is also enriched on strong boundaries of CIDs and a deletion of Mediator Med1 subunit modifies chromatin compaction (Chereji et al., 2017; Hsieh *et al*., 2015). Consistent with physical interactions between Mediator and RSC complexes, mass spectrometry analysis of coimmunoprecipitated proteins showed that chromatin-bound Mediator is associated with RSC subunits (Chereji *et al*., 2017). Future studies will help to understand how the Mediator link to RSC can have an impact on higher-order genome organization.

Previously, we showed that RSC interacts with RNA polymerases and that the loss of this interaction alters the promoter chromatin structure of several RSC-regulated genes, leading to impaired transcription (Soutourina et al., 2006). These results have led us to propose that RSC might act as a coactivator independently of its catalytic activity in chromatin remodeling. Further studies will be needed to understand the exact relationships between chromatin remodeling activity of RSC and its effect on transcription independent of its catalytic activity. In the light of Mediator and SWI-SNF remodelers PBAF/BAF/ncBAF conservation from yeast to humans, the functional cooperation between these complexes is likely to exist in all eukaryotes. The conservation of Mediator connection to chromatin remodelers might give insights into our understanding of human diseases such as many cancers involving Mediator and PBAF/BAF/ncBAF subunits.

## Acknowledgements

We thank Charles Boone, Jenny Wu, Claes Gustafsson and Vincent Geli for strains, the High-throughput sequencing facility of I2BC for its sequencing and bioinformatics expertise, the SPI (CEA/Saclay) for monoclonal antibodies, Matthieu Gérard, Pascale Lesage and Adriana Alberti for fruitful discussions. This work was supported by Fondation ARC [PGA1 RF20170205342]; Comité Ile-de-France - La Ligue Nationale Contre le Cancer. K.A. was supported by a PhD training contact from the French Ministry of Higher Education and Research. L.Z. was supported by a PhD training contract from the CEA NUMERICS program, which has received funding from European Union’s Horizon 2020 research and innovation program under the Marie Sklodowska-Curie grant agreement No 800945.

## Author contributions

JS and MW designed the experiments. KA, NGA and VMF performed the experiments. KA and LZ performed data analysis with supervision of CDW and AG. Supervision and project coordination was done by JS. JS and KA wrote the original draft. Review and editing were done by JS, KA, CDW, AG, MW and LZ.

## Declaration of interests

The authors declare no competing interests.

## Materials and Methods

### Strains

All *Saccharomyces cerevisiae* strains used in this study can be found in **Supplementary Table S1**.

*med17* mutant strains were previously described (Eyboulet *et al*., 2013; Eyboulet *et al*., 2015). *rsc8* and *sth1* mutant strains were obtained from Charles Boone (Li *et al*., 2011). *rsc8* and *sth1* mutations associated with *KanMX* cassette were then transferred to YPH499 background by the standard one-step methods. Med14-Myc, Rsc8-Myc, Sth1-Myc strains carrying C-terminal Myc-tagged version of Med14, Rsc8 and Sth1 subunits were constructed by inserting Myc epitopes followed by either *KanMX, His3MX6* or *LEU2* markers using the standard one-step methods. Prior to ChIP-qPCR, ChIP-seq, MNase-seq and CoIP experiments, Mediator mutant strains were shifted for 45 min at 37°C, *rsc8-ts* strains were shifted for 6h at 37°C, and *sth1-ts* strains were shifted for 2h30 at 37°C. Spotting assays were done as previously described (Georges *et al*., 2019).

### Yeast two-hybrid assay

Haploid strains (Y187 and Y190) were transformed with constructs expressing the desired gene fusion with Gal4 DNA-binding domain (*TRP* auxotrophic marker) or Gal4 activating domain (*LEU* auxotrophic marker). Clones growing on –*trp* or –*leu* medium were selected and the presence of the plasmid was verified by PCR.

Haploid strains growing on –*trp* and –*leu* agar plates were scraped and resuspended in sterile water. The strains were then mated together by spotting 2.5 µl of each suspension on a YPD plate and incubating overnight at 30°C. Patches were replica-plated on –*trp –leu* plates and incubated for 3 days at 30°C. Diploid strains were scraped and resuspended in synthetic defined (SD) medium supplemented with 40 mg/l Adenine (SD+2A). Optical density at 600 nm of the suspension was measured and the latter was diluted to obtain an OD_600_ of 0.1. 10 µl of this dilution were then spotted on agar plates containing SD+2A medium supplemented with 10, 25 or 50 mM 3-amino-1,2,4-triazole (3AT), and incubated for 3 days at 30°C.

When indicated, X-gal staining was done as follows: for one 120 mm square petri dish, 10 ml 1% Agarose in water was mixed with 10 ml phosphate buffer (made from 6.15 ml 1M K_2_HPO_4_ and 3.85 ml 1M KH_2_PO_4_ aqueous solutions), both prewarmed at 50°C. 1.2 ml N,N-dimethylformamide (DMF), 0.2 ml 10% sodium dodecyl sulphate (SDS) solution (in ddH_2_O) and 0.2 ml of 4% X-Gal solution (in DMF, kept at −20°C) were successively added to the mix. The mix was then poured onto yeast spots. As soon as solidified, the plate was incubated at 30°C for 24h. Pictures were taken using an office scanner.

### Coimmunoprecipitation

A total of 100 ml exponentially growing cells were collected by centrifugation, washed and cell lysis was performed by bead-beating for 30 min at 4°C in lysis buffer (20% Glycerol, 50 mM Hepes-KOH pH 7.5, 100 mM NaCl, 0.5 mM ethylenediaminetetraacetic acid (EDTA), 0.05% NP-40, 1 mM dithiothreitol (DTT), 1 mM phenylmethylsulfonide fluoride (PMSF), Complete protease inhibitor cocktail (Roche)) as described previously (Soutourina *et al*., 2006). Protein concentration was measured using Bradford method, taking bovine serum albumin (BSA) as reference.

Protein extracts were used for immunoprecipitation as follows: 50 µl Dynabeads pan-mouse IgG were washed three times in cold phosphate-buffered saline (PBS) containing 0.1% BSA, and incubated 30 min at 4°C on rotary wheel in the same solution. The beads were then incubated 1 h with the anti-Myc antibody (9E10) at 4°C (Eppendorf ThermoMixer, 1,300 rpm), washed three times in PBS/0.1 % BSA and two times in lysis buffer. A total of 1.5 mg of proteins were added to beads and lysis buffer was added to have the same final volume in all samples (at least 50 µl). Beads and proteins were incubated for 3 hours at 4°C (Eppendorf ThermoMixer, 1,300 rpm). Beads were then washed four times with lysis buffer and recovered in 40 µl of laemmli buffer (50 mM Tris-Cl pH 6.8, 0.5 % sodium dodecyl sulfate (SDS), 2.5 % Glycerol, 1.25 % β-mercaptoethanol, 0.0125 % Bromophenol blue). Proteins were eluted at 95°C for 10 min and stored at −80°C.

Prior to SDS-polyacrylamide gel electrophoresis, samples were incubated at 85°C for 5 minutes. Separation was done on 8% bis-acrylamide gels in Tris-Glycine-SDS buffer and proteins were transferred on Amersham Protran 0.2 µm NC membranes (GE Healthcare) for western blotting. Membranes were bathed in PBS supplemented with 0.05% Tween 80 (PBS-T) with 5% Milk for 1 h at room temperature, then incubated overnight at 4°C with either anti-Myc (9E10) or anti-Med14 (Eyboulet *et al*., 2015) antibodies. After three washes in PBS-T with 2% Milk the membranes were incubated with secondary antibodies (HRP-anti Rabbit/Mouse-IgG (H+L), Promega) for 1 hour at room temperature in PBS-T. After three more washes in PBS-T, detection was carried out using Amersham ECL-Prime reagents (GE Healthcare). Imaging was done using a Fusion FX7 imaging system.

The relative intensity of immune staining was quantified using ImageJ software. The intensity of immune staining for coimmunoprecipitated protein signals relative to the WT was normalized against immunoprecipitation and input signals. The mean values and standard deviation (indicated by error bars) of three independent experiments were calculated.

### Chromatin immunoprecipitation

Cells were grown exponentially to 0.6 OD_600_ and cross-linked with 1% formaldehyde for 10 min. Cells were lysed by bead-beating in FA/SDS buffer (50 mM Hepes-KOH pH 7.5, 150 mM NaCl, 1 mM EDTA, 1% Triton, 0.1% sodium deoxycholate, 0.1% SDS) supplemented with PMSF for 30 min at 4°C. Chromatin was recovered after a centrifuge step (13,400 rcf, 20 min, 4°C) and was subjected to sonication on a S220 focused-ultrasonicator (Covaris) (180s ON – 30s OFF – 180s ON, 150W pulses, duty factor 10). Sonicated chromatin was recovered after centrifuge (9,300 rcf, 30 min, 4°C) and stored at −80°C. The sizes of the DNA fragments were checked by migrating the samples on agarose.

Immunoprecipitation, DNA purification and library preparation were done on IP-Star® Compact Automated System (Diagenode Cat# B03000002).

IP was done as previously described (Georges *et al*., 2019). Anti-Myc (9E10), anti-Pol II (8WG16) and anti-H3 (Merck, 07-690) antibodies were used. Spike-in *S. pombe* chromatin was added to the different samples at the IP step (**Supplementary Table S2**). Crosslink reversion, RNase treatment and DNA purification were done as previously described (Georges *et al*., 2019), except for the RNase treatment duration which was 1h for this current study.

qPCR and libraries preparations were done as previously described (Georges *et al*., 2019). qPCR primers can be found in **Supplementary Table S3**.

### MNase treatment

Cells were grown exponentially to 0.6 OD600 and cross-linked with 1% formaldehyde for 10 min. They were then washed with ST buffer (10 mM Tris-Cl pH 7.5, 100 mM NaCl) and frozen. Cells were lysed by bead-beating in FA lysis buffer (50 mM Hepes-KOH pH 8, 150 mM NaCl, 0.5 mM EDTA, 1% Triton, 0.1% sodium deoxycholate) supplemented with PMSF for 30 minutes at 4°C. Chromatin was recovered after a centrifuge step (16,000 rcf, 10 min, 4°C) in MNase buffer (20 mM Tris-HCl pH 7.5, 0.34 M sucrose, 15 mM KCl, 60 mM NaCl, 3 mM CaCl_2_) and was subjected to mild MNase (NEB, M0247S) treatment (20 min, 37°C on thermomixer). Reaction was stopped by addition of 10 mM EDTA on ice. Digested chromatin was recovered after centrifugation (16,000 rcf, 10 min, 4°C), and the remaining pellet was mildly sonicated (20” ON + 40” OFF, 4 cycles, mild), after a centrifugation step under similar conditions the chromatin samples were recovered at −80°C. DNA was extracted with PCR purification kit (Qiagen). The sizes of the DNA fragments were checked by migrating the samples on both hand-made agarose and BioAanalyzer (Agilent) after pronase treatment. MNAse digestion profiles were comparable between different samples and contained a major mono-nucleosome peak with clearly visible di- and tri-nucleosome peaks.

Library preparations were done as previously described (Georges *et al*., 2019), except for the sizing step that was hand-made following the AMPure protocol (Beckman) to remove the free sequencing primers only. Library profiles were checked by migrating them on BioAanalyzer (Agilent).

### Sequencing data analysis

Sequencing data were analyzed using the following procedure. Reads were first trimmed with cutadapt (v1.12, http://dx.doi.org/10.14806/ej.17.1.200), then mapped on a *S. cerevisiae* + *S. pombe* concatenated genome (University of California at Santa Cruz [UCSC] versions sacCer3 and ASM284v2, respectively) using bowtie2 (v2.3.4.3, doi: 10.1038/nmeth.1923). Files were converted using samtools (v1.9, doi: 10.1093/bioinformatics/btp352) and deeptools (v3.5.0, doi: 10.1093/nar/gkw257). After bam conversion, *S. cerevisiae* alignments were separated from *S. pombe* alignments and the spike-in normalization factors were determined. Read count were then normalized in reads per million of mapped reads (RPM) and spike-in normalization factors. MNase-seq experiments were only normalized in RPM. The number of mapped reads for each sequencing experiment and normalization coefficients are indicated in **Supplementary Table S4**. For MNase-seq experiments, the option --MNase of bamCoverage was used so that only the mononucleosome fragments were kept (fragments shorter than 130 bp and longer than 200 bp were removed from analysis). Input DNA and DNA from ChIP with a non-tagged strain were used as negative controls.

The transcribed regions were determined using the TSS and TES (transcription end sites) of mRNA genes taken from (Malabat et al., 2015; Pelechano et al., 2013) (n=5337). The lengths of the nucleosome-depleted regions of promoters were calculated from the WT MNase-seq data as the distance between the −1 and +1 nucleosomes, which were defined as the closer nucleosomes upstream and downstream from the TSS, respectively (https://doi.org/10.5281/zenodo.6669875). Peak calling was done on Med17, Rsc8 and Sth1 experiments using MACS2 (v2.2.7.1). The peaks were filtered (fold-change >2.5 and *P*-value < 1e-10), and as we wanted to focus on protein-coding gene promoters, we kept those that were found on intergenic regions associated to protein-coding genes and <1 kb away from TSSs. We excluded intergenic regions related to replication origins, centromeres, long-terminal repeats, Pol III and Pol I-transcribed genes, retrotransposons, small nucleolar RNAs, small nuclear RNAs and telomeres. For some analysis, peaks were sorted using bedtools (v2.30.0) to define those that overlap (shared) or not (unique) between two data sets. All correlations were calculated with Spearman method (Hollander, 1973). All figures were prepared using R packages (v3.5.1, https://www.R-project.org/). Specifically, heatmap with sorted nucleosome-depleted regions was done with R using the package ComplexHeatmap.

### Data availability

The ChIP-seq and MNase-seq data generated in this study have been submitted to ArrayExpress (https://www.ebi.ac.uk/arrayexpress/) under the accession number E-MTAB-12198.

## Notes

### Competing Interest Statement

The authors have declared no competing interest.

