## Supplementary Data for "Functional interplay between Mediator and RSC chromatin remodeling complex controls nucleosome-depleted region maintenance at promoters"

**Supplementary Table S1. Yeast strains used in the study.**

| Strain name | *Genotype* | Collection # | Reference |
|---|---|---|---|
| YPH499 | *MATa ade2-1 lys2-801 ura3-52 trp1-Δ63 his3-Δ200 leu2Δ* | 97 | (Sikorski & Hieter, 1989)* |
| med17Δ/MED17 TRP Rsc8-Myc | *MATa ura3-52 his3-Δ200 ade2-101uaa trp1-Δ63 lys2-801uag leu2-Δ1 Δmed17::kan::ADE2 MED17:CEN:TRP1 RSC8:13MYC:KANMX* | 7977 | This work |
| med17Δ/med17-140 (Q444P M442L) TRP Rsc8-Myc | *MATa ura3-52 his3-Δ200 ade2-101uaa trp1-Δ63 lys2-801uag leu2-Δ1 Δmed17::kan::ADE2 med17-140:CEN:TRP1 RSC8:13MYC:KANMX* | 7978 | This work |
| med17Δ/med17-444 (Q444P) TRP Rsc8-Myc | *MATa ura3-52 his3-Δ200 ade2-101uaa trp1-Δ63 lys2-801uag leu2-Δ1 Δmed17::kan::ADE2 med17-444:CEN:TRP1 RSC8:13MYC:KANMX* | 7979 | This work |
| med17Δ/MED17 TRP Sth1-Myc | *MATa ura3-52 his3-Δ200 ade2-101uaa trp1-Δ63 lys2-801uag leu2-Δ1 Δmed17::kan::ADE2 MED17:CEN:TRP1 STH1:13MYC:KANMX* | 7982 | This work |
| med17Δ/med17-140 (Q444P M442L) TRP Sth1-Myc | *MATa ura3-52 his3-Δ200 ade2-101uaa trp1-Δ63 lys2-801uag leu2-Δ1 Δmed17::kan::ADE2 med17-140:CEN:TRP1 STH1:13MYC:KANMX* | 7983 | This work |
| med17Δ/med17-444 (Q444P) TRP Sth1-Myc | *MATa ura3-52 his3-Δ200 ade2-101uaa trp1-Δ63 lys2-801uag leu2-Δ1 Δmed17::kan::ADE2 med17-444:CEN:TRP1 STH1:13MYC:KANMX* | 7984 | This work |
| med17Δ/MED17 TRP Med14-Myc | *MATa ura3-52 his3-D200 ade2-101uaa trp1-D63 lys2-801uag leu2-D1 Δmed17::kan::ADE2 MED14Myc HIS CEN MED17 TRP* | 7995 | This work |
| med17Δ/med17-140 (Q444P M442L) TRP Med14-Myc | *MATa ura3-52 his3-D200 ade2-101uaa trp1-D63 lys2-801uag leu2-D1 Δmed17::kan::ADE2 MED14Myc HIS CEN med17-140 TRP* | 7996 | This work |
| med17Δ/med17-444 (Q444P) TRP Med14-Myc | *MATa ura3-52 his3-D200 ade2-101uaa trp1-D63 lys2-801uag leu2-D1 Δmed17::kan::ADE2 MED14Myc HIS CEN med17-444 TRP* | 7998 | This work |

| | | | |
|---|---|---|---|
| med17Δ/MED17 URA3 Med14-Myc | *MATa ura3-52 his3-D200 ade2-101uaa trp1-D63 lys2-801uag leu2-D1 Δmed17::kan::ADE2 MED17 CEN URA3 MED14::13myc::HIS3MX6* | 7987 | This work |
| med17Δ/MED17 URA3 rsc8-ts (G326D I417F K446M) Med14-Myc | *MATa ura3-52 his3-D200 ade2-101uaa trp1-D63 lys2-801uag leu2-D1 Δmed17::kan::ADE2 MED17 CEN URA3 MED14::13myc::HIS3MX6 rsc8-ts21 (G326D, I417F, K446M)::KanR* | 8026 | This work |
| med17Δ/MED17 URA3 sth1-ts (P646L F655L D657G N1077S F1125L) Med14-Myc | *MATa ura3-52 his3-D200 ade2-101uaa trp1-D63 lys2-801uag leu2-D1 Δmed17::kan::ADE2 MED17 CEN URA3 MED14::13myc::HIS3MX6 sth1-ts (P646L, F655L, D657G ,N1077S,F1125L)* | 8034 | This work |
| med17Δ/MED17 URA3 Rsc8-Myc | *MATa ura3-52 his3-D200 ade2-101uaa trp1-D63 lys2-801uag leu2-D1 Δmed17::kan::ADE2 MED17 CEN URA3 RSC8::13MYC::KAN::LEU2* | 7989 | This work |
| med17Δ/MED17 URA3 sth1-ts (P646L F655L D657G N1077S F1125L) Rsc8-Myc | *MATa ura3-52 his3-D200 ade2-101uaa trp1-D63 lys2-801uag leu2-D1 Δmed17::kan::ADE2 MED17 CEN URA3 RSC8::13MYC::KAN::LEU2 sth1-ts( P646L F655L , D657G ,N1077S,F1125L)* | 8069 | This work |
| med17Δ/MED17 URA3 Sth1-Myc | *MATa ura3-52 his3-D200 ade2-101uaa trp1-D63 lys2-801uag leu2-D1 Δmed17::kan::ADE2 MED17 CEN URA3 STH1::13MYC::KAN::LEU2* | 7988 | This work |
| med17Δ/MED17 URA3 rsc8-ts (G326D I417F K446M) Sth1-Myc | *MATa ura3-52 his3-D200 ade2-101uaa trp1-D63 lys2-801uag leu2-D1 Dmed17::kan::ADE2 MED17 CEN URA3 STH1::13MYC::KAN::LEU2 rsc8-ts21(G326D, I417F, K446M)::KanR* | 8068 | This work |

*Sikorski RS, Hieter P. 1989. A system of shuttle vectors and yeast host strains designed for efficient manipulation of DNA in *Saccharomyces cerevisiae*. Genetics 122: 19-27.

#### Supplementary Table S2. Spike-in parameters for ChIP-seq.

| ***S. cerevisiae* strain** | **IP** | **Spike-in fraction (%)** | ***S. pombe* strain** |
|---|---|---|---|
| *med17* mutants | Pol II | 5 | Rif1-Myc |
| *med17* mutants | Rsc8-Myc | 10 | Rif1-Myc |
| *med17* mutants | Med14-Myc | 10 | Med4-Myc |
| *med17* mutants | Sth1-Myc | 20 | Rif1-Myc |
| Non-tagged | Myc | 10 | Rif1-Myc |
| *rsc8* mutant | Med14-Myc | 20 | Med7-Myc |
| *rsc8* mutant | Pol II | 10 | Med7-Myc |
| *rsc8* mutant | Sth1-Myc | 20 | Med7-Myc |
| *sth1* mutant | Med14-Myc | 20 | Med7-Myc |
| *sth1* mutant | Pol II | 10 | Med7-Myc |
| *sth1* mutant | Rsc8-Myc | 20 | Med7-Myc |

**Supplementary Table S3. Oligonucleotides used in the study.**

| qPCR primers | SEQUENCE | Primer pair |
|---|---|---|
| **GAL1 ORF** | GAL1-O1 / O2 | AAAGAAACTTGCACCGGAAA/ GGCCCATATTCGCTTTAACA |
| **IGV** | IGV-3 / 4 | TTCATTTGCAATTTGCAGTTCA/ CAGCCAGGAAGAATCTCACAA |
| **PYK1 UAS** | PYK1-P1 / P2 | CGCACCGTCACAAAGTGTT/ TGGGAAGGAAAGGAAATCAC |
| **PYK1 CP** | PYK1-P3 / P4 | CCTTTCCTTCCCATATGATGC/ ACTTTGAAAGGGGACCATGA |
| **PYK1 ORF** | PYK1-O1 / O2 | ATGGTTGCCAGAGGTGACTT/ TCTGGTTGGTCTTGGGTTGT |
| **PMA1 UAS** | PMA1-P3 / P4 | AACAAACCCGGTCTCGAAG/ GAAGTGCCGCATTAGGAAAT |
| **PMA1 CP** | PMA1-378f / 235r | GATGGTGGGTACCGCTTATG/ TTGGTGTTATAGGAAAGAAAGAGAAAA |
| **PMA1 ORF** | PMA1-O1 / O2 | GGTTTTGGTCATTGCCACTT/ ACGGCCATAGTGGTGGTAAC |
| **SNQ2 UAS** | SNQ2-P1N / P2N | TCACCCCATTTGGAAAAGAA/ TAGTCATATGTGCGCGGAAC |
| **SNQ2 CP** | SNQ2-P3N / P4N | GCCCATTTCCGTTTAAATCC/ GAGGGAGGGGCACTACTCAT |
| **SNQ2 ORF** | SNQ2-O1 / O2 | GTTCAGAGGCGATCAATGGT/ ATCGAAGGCAATTGGATCAT |

**Supplementary Table S4. Number of mapped reads and normalization coefficient of each sequencing sample.**

| Sample | Number of mapped reads on *S. cerevisiae* | Normalization factor |
|---|---|---|
| Input_Rsc8-Myc_WT | 22977800 | 1 |
| Pol II_Rsc8-Myc_WT | 16516026 | 1 |
| Pol II_Rsc8-Myc_med17-140 | 9995666 | 0.39 |
| Pol II_Rsc8-Myc_med17-444 | 12849238 | 0.49 |
| Rsc8_Rsc8-Myc_WT | 10784068 | 1 |
| Rsc8_Rsc8-Myc_med17-140 | 10747200 | 0.67 |
| Rsc8_Rsc8-Myc_med17-444 | 9942148 | 1.69 |
| Med14_Med14-Myc_WT | 3993990 | 1 |
| Med14_Med14-Myc_med17-140 | 3186594 | 0.7 |
| Med14_Med14-Myc_med17-444 | 2785480 | 1.12 |
| Sth1_Sth1-Myc_WT | 17239958 | 1 |
| Sth1_Sth1-Myc_med17-140 | 16445980 | 0.83 |
| Sth1_Sth1-Myc_med17-444 | 18320564 | 0.67 |
| Med14_Med14-Myc_WT6h | 10865416 | 1 |
| Med14_Med14-Myc_rsc8-ts | 5973146 | 0.91 |
| Pol II_Med14-Myc_WT6h | 19463068 | 1 |
| Pol II_Med14-Myc_rsc8-ts | 13954246 | 0.39 |
| Sth1_Sth1-Myc_WT6h | 19800880 | 1 |
| Sth1_Sth1-Myc_rsc8-ts | 5820932 | 0.84 |
| Med14_Med14-Myc_WT2h | 13006740 | 1 |

| | | |
|---|---|---|
| Med14_Med14-Myc_sth1-ts | 3424764 | 1.39 |
| Pol II_Med14-Myc_WT2h | 21245530 | 1 |
| Pol II_Med14-Myc_sth1-ts | 19081108 | 0.38 |
| Rsc8_Rsc8-Myc_WT2h | 23298254 | 1 |
| Rsc8_Rsc8-Myc_sth1-ts | 20475408 | 0.98 |
| IP-Myc_non-tagged_strain_WT | 2167402 | 1 |
| MNase_Rsc8-Myc_WT | 24773602 | 1 |
| MNase_Rsc8-Myc_med17-140 | 16600296 | 1 |
| MNase_Rsc8-Myc_med17-444 | 16341674 | 1 |
| MNase_Med14-Myc_WT6 | 20707292 | 1 |
| MNase_Med14-Myc_rsc8-ts | 18523968 | 1 |
| MNase_Med14-Myc_WT2 | 22068258 | 1 |
| MNase_Med14-Myc_sth1-ts | 20485700 | 1 |

**Supplementary Figure Legends**

**Supplementary Figure S1.** Med21 interacts with a fragment of Rsc3. Gal4 DNA-binding domain ($G_{BD}$), alone or in fusion with full length Med21, Med11 or Med7, was co-expressed with Gal4 activation domain ($G_{AD}$) alone or in fusion with full length Rsc30, partially truncated Rsc30 (Rsc30F: amino acids 644-883) or partially truncated Rsc3 (Rsc3F: amino acids 588-695). Yeast cells were spotted on SD+2A medium supplemented with 25 mM 3-AT, grown for 3 days (left panel) and then stained with X-Gal for 24h (right panel).

**Supplementary Figure S2. Mediator and RSC colocalize on NDRs within promoter regions.** (**A**) Average tag densities of Med14, Sth1, Rsc8, Pol II and IP Myc in a non-tagged strain on *GIC2, FUI1, MAL33* and *SNQ2* genes. (**B**) Venn diagrams illustrating the number of Med14 peaks that overlap or not with Sth1 and Rsc8 peaks. (**C**) Heat maps and average profiles of Med14-Myc, Sth1-Myc and Rsc8-Myc centered on Rsc8 and Sth1 peaks within intergenic regions, -500 bp to +500 bp. Signals are expressed as the normalized read counts. (**D**) Spearman correlation coefficients of ChIP-seq data between Med14, Rsc8 and Sth1 on intergenic regions. The colors correspond to the scale for Spearman correlation coefficients indicated on the bottom. (**E**) Sth1, Rsc8, Med14 and MNase profiles centered on TSSs associated to Rsc8 and

Sth1 peaks, -500 bp to +500 bp. The profiles were normalized for comparison by rescaling their maximum values to 1.

**Supplementary Figure S3. Effects of *rsc* mutations on Pol II, Med14 and RSC chromatin occupancy.** (**A**) Boxplots showing the Pol II enrichment in WT, *rsc8-ts* and *sth1-ts* strains. The three asterisks represent a significant difference between the WT and the mutant at P-value < 0.001 in a Wilcoxon test. (**B**) Top panel: heatmaps and average profiles of Med14-Myc and Sth1-Myc ChIP-seq signals centered on Sth1 WT peaks (n=763, -500 bp to +500 bp) in WT or *rsc8-ts*. Bottom panel: heat maps and average profiles of Med14-Myc and Rsc8-Myc ChIP-seq signals in WT or *sth1-ts* centered on Rsc8 WT peaks (n=464), -500 bp to +500 bp. Negative control was obtained from a Myc IP in a non-tagged strain (NT). Signals are expressed as the normalized read counts.

**Supplementary Figure S4. Effects of *rsc* mutations on Pol II, Med14 and RSC chromatin occupancy.** Quantitative ChIP analysis of Pol II, Mediator, Sth1 and Rsc8 occupancies following a 6-hour or 2.5-hour shift at 37°C (for *rsc8-ts* and *sth1-ts*, respectively), after reaching exponential phase at 25°C. Quantitative PCR (qPCR) was performed on precipitated DNA, using primer pairs designed to amplify either regions in open reading frames (ORF) or upstream activating sequences (UAS). Relative quantity of an amplicon was determined by comparing the obtained Ct to a standard curve made on the same qPCR plate. Quantities were reported to qPCR performed on input DNA and are expressed as a percentage. *GAL1* ORF amplicon was used as a negative control. The indicated value is the mean of three biological replicates, and error bars represents the standard deviation.

**Supplementary Figure S5. Effect of Mediator mutations on NDRs and protein occupancies.** (**A**) Boxplots showing the Pol II enrichment in WT, *med17-140* and *med17-444* strains. The three asterisks represent a significant difference between the WT and the mutant at P-value < 0.001 in a Wilcoxon test. (**B**) Venn diagrams illustrating the number of Med14 peaks that overlap or not with Sth1 (left) and Rsc8 (right) peaks in *med17-140*. (**C**) Heat maps and average profiles of Med14-Myc, Sth1-Myc and Rsc8-Myc centered on Sth1 peaks within intergenic regions, -500 bp to +500 bp, in *med17-140* (left panel) or *med17-444* (right panel). Signals are expressed as the normalized read counts. (**D**) Spearman correlation coefficients of ChIP-seq data between Med14, Rsc8 and Sth1 on intergenic regions in *med17-140* (left panel) and *med17-444* (right panel). The colors correspond to the scale for Spearman correlation coefficients indicated on the bottom. (**E**) Boxplots showing the Sth1 (top left), Rsc8 (top right) and Med14 (bottom left) enrichments in WT, *med17-140* and *med17-444* strains. The three asterisks represent a significant difference between the WT and the mutant at P-value < 0.001 in a Wilcoxon test. Negative control was obtained from a Myc IP in a non-tagged strain (NT). (**F**) Heat maps and average MNase profiles in WT, *med17-140* and *med17-444* strains centered on Rsc8 (left panels) or Med14 (right panels) peaks within intergenic regions (n=337 and 313, respectively), -500 bp to +500 bp. Signals are expressed as the normalized read counts.

**Supplementary Figure S6. Effects of *med17* mutations on Pol II, Med14 and RSC chromatin occupancy.** Quantitative ChIP analysis of Pol II, Med14, Sth1 and Rsc8 occupancies following a 45-min shift at 37°C, after reaching exponential phase at 30°C. Quantitative PCR (qPCR) was performed on precipitated DNA, using primer pairs designed to amplify either regions in open reading frames (ORF), upstream activating sequences (UAS) or core promoters (CP). Relative quantity of an amplicon was determined by comparing the obtained Ct to a standard curve made on the same qPCR plate. Quantities were reported to qPCR performed on input DNA and are expressed as a percentage. *GAL1* ORF and *IGV* (intergenic region on chromosome V) amplicons were used as negative controls. The indicated value is the mean of three biological replicates, and error bars represents the standard deviation.

**Supplementary Figure S7. H3 occupancy is increased on promoters in *med17-140*.** (**A**) Quantitative ChIP analysis of H3 occupancy following a 45-min shift at 37°C, after reaching exponential phase at 30°C. Quantitative PCR (qPCR) was performed on precipitated DNA, using primer pairs designed to amplify either regions in open reading frames (ORF), upstream activating sequences (UAS) or core promoters (CP). Relative quantity of an amplicon was determined by comparing the obtained Ct to a standard curve made on the same qPCR plate. Quantities were reported to qPCR performed on input DNA and are expressed as a percentage. *GAL1* ORF and *IGV* (intergenic region on chromosome V) amplicons were used as non-transcribed controls. The indicated value is the mean of three biological replicates, and error bars represents the standard deviation. The two asterisks represent a significant difference between the WT and the mutant at P-value < 0.01 in a Student t-test, while the single asterisk represents a significant difference at P-value < 0.05. (**B**) Examples of MNase-seq profiles in WT and *med17* mutants on selected UASs. Average tag density of Med14 ChIP-seq in WT is also shown in orange. Positions of qPCR primers for UASs are indicated by red boxes.

**Supplementary Figure S8. NDRs are disturbed in *med17-140* on promoters enriched by both Mediator and RSC.** (**A**) Heat maps and average MNase profiles from the WT and *med17-140* strains centered on *med17-140* Rsc8 peaks within intergenic regions that overlap (left panel) or not (right panel) with Med14 peaks (n=310 and 531, respectively), -500 bp to +500 bp. Signals are expressed as the normalized read counts. (**B**) Average MNase profiles in WT and *med17-140* centered on TSSs associated to Rsc8/Med14-shared peaks (n=310) or Rsc8 only (n=531), -500 bp to +500 bp. The profiles were normalized for comparison by rescaling their maximum values to 1. (**C**) Boxplots showing Rsc8 and Med14 enrichments (left and right panels, respectively) in WT and *med17-140* on *med17-140* Sth1 peaks that overlap (termed shared) or not (termed unique) with Med14 peaks. (**D**) Boxplots showing Sth1, Rsc8 and Med14 enrichments (left, middle and right panels, respectively) in WT and *med17-140* on *med17-140* Rsc8 peaks that overlap (termed shared) or not (termed unique) with Med14 peaks. (**E**) Boxplots showing the NDR lengths on promoters enriched by both Rsc8 and Med14 (termed shared) or Rsc8 only (termed unique), in comparison to all NDRs. The three asterisks represent a significant difference between the Rsc8/Med14-associated NDRs and all NDRs or Rsc8-associated NDRs at P-value < 0.001 in a Wilcoxon test. (**F**) Density plot showing NDR length distribution for promoter NDRs associated to Med14 and Sth1 (left) or Sth1 only (right).

**Supplementary Figure S9. Genes associated to shared Mediator/RSC peaks are more occupied by Pol II than genes associated to unique RSC peaks.** (**A**) Boxplots showing Pol II enrichment in WT and *med17-140* on *med17-140* Sth1 and Rsc8 peaks that overlap (termed shared) or not (termed unique) with Med14 peaks. (**B**) *med17-140*/WT ratios of Pol II occupancy on coding regions associated to Sth1 and Rsc8 peaks (left and right panels, respectively), shared between RSC and Mediator or unique. Ratios were calculated from the mean Pol II tag densities on coding regions.

### Supplementary Figure S1

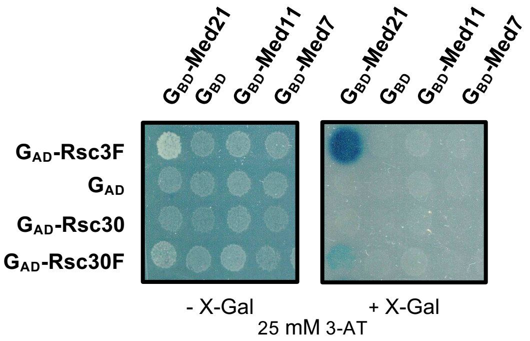

### Supplementary Figure S2

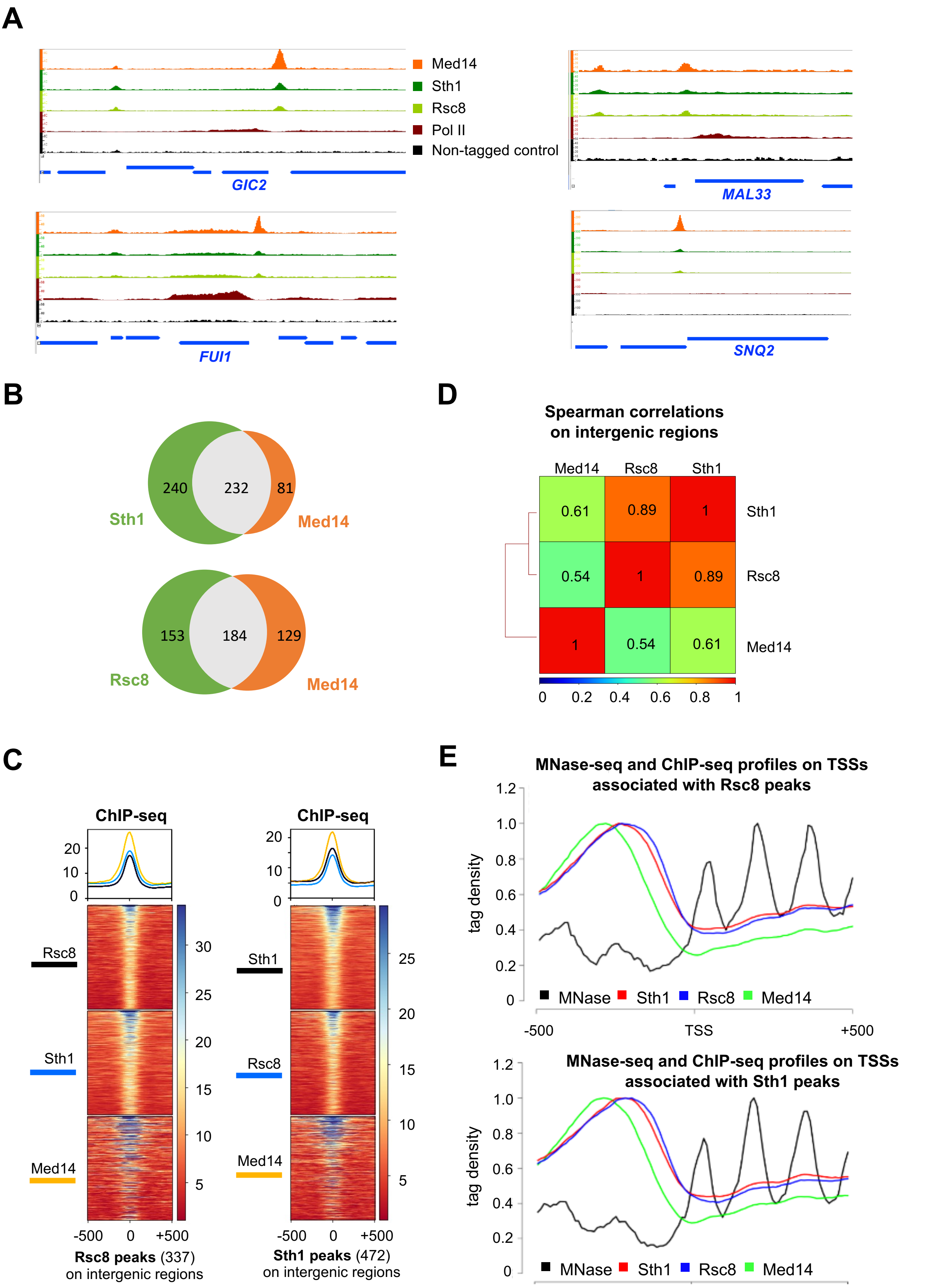

### Supplementary Figure S3

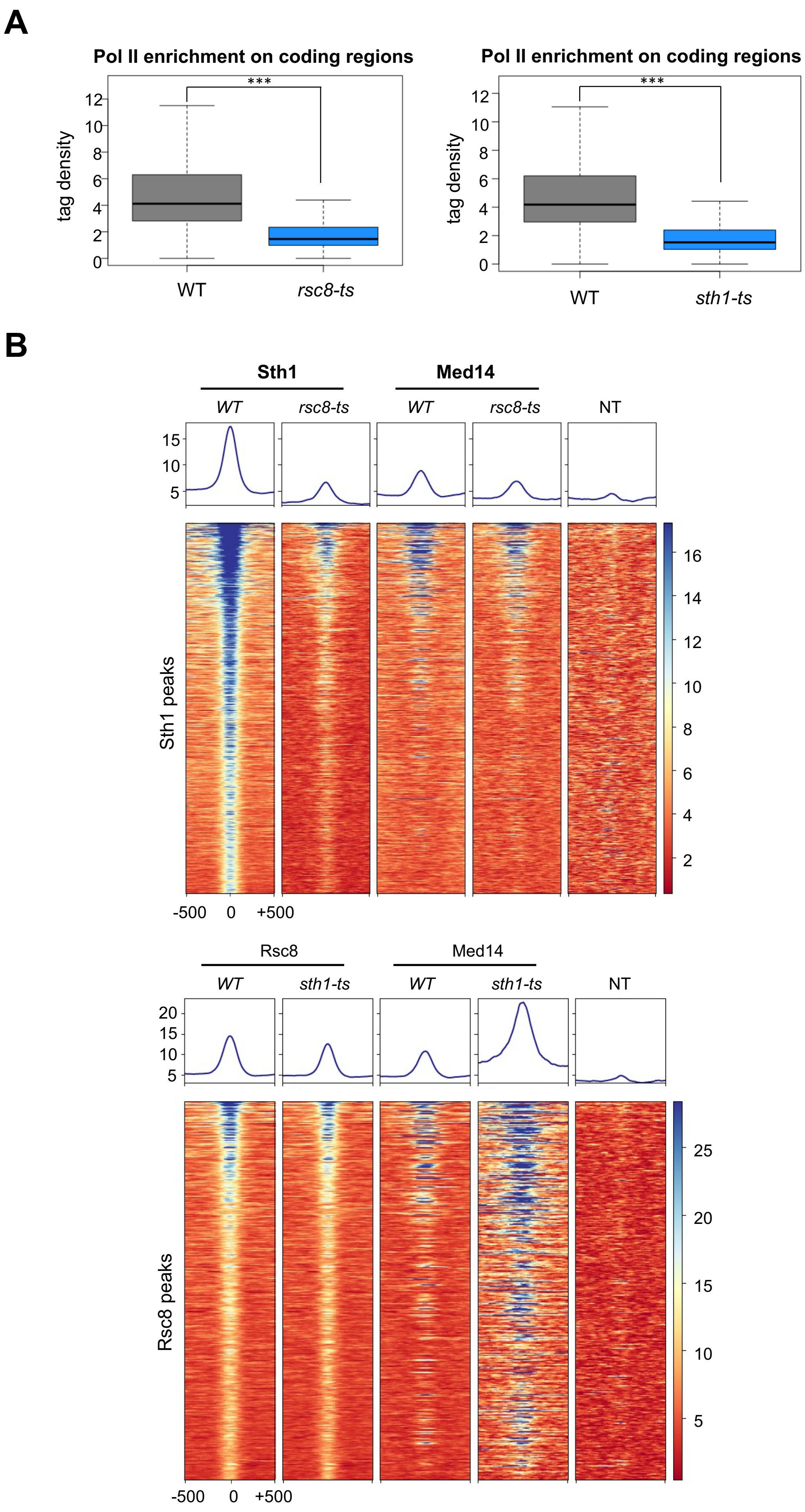

### Supplementary Figure S4

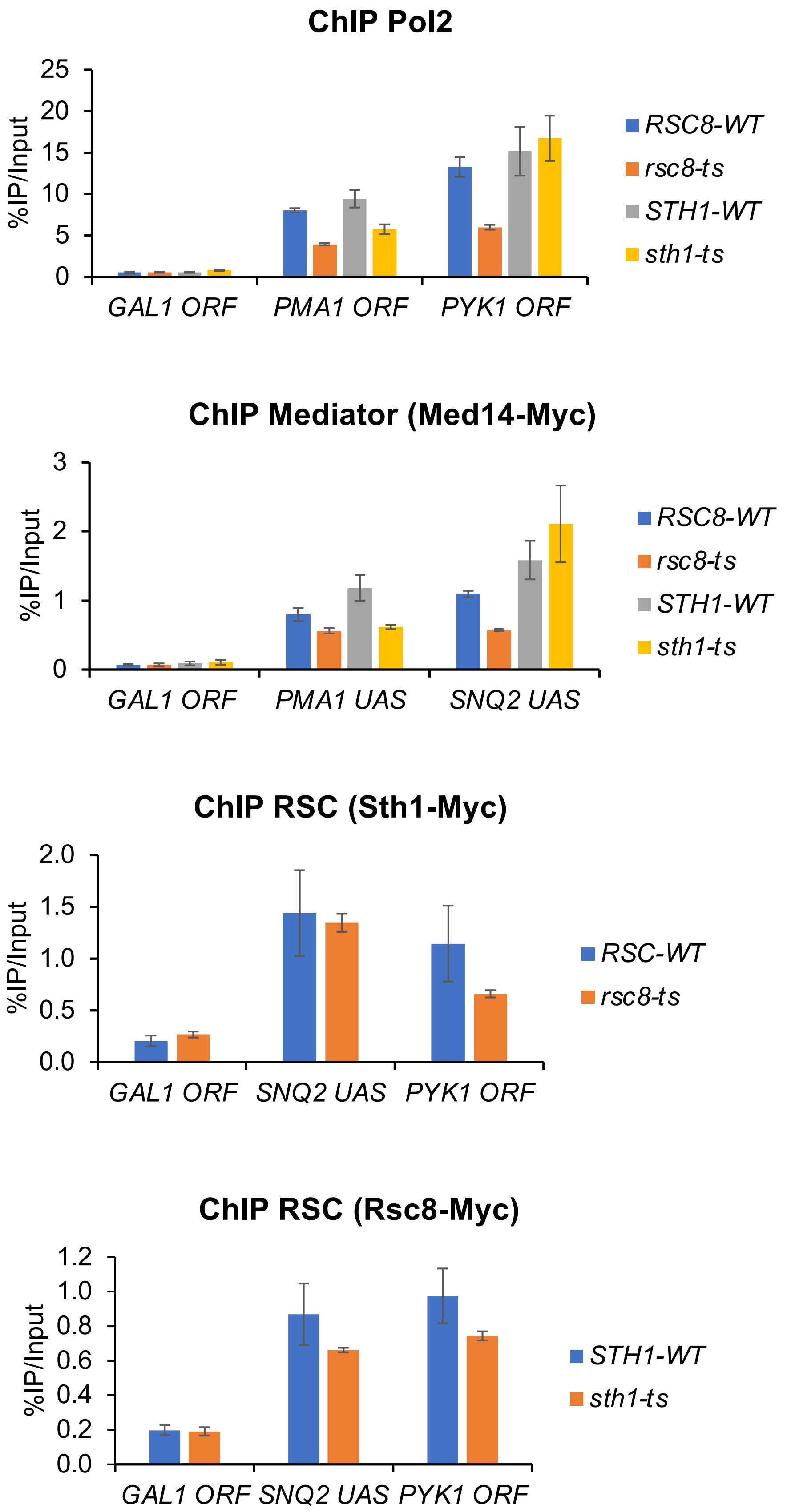

### Supplementary Figure S5

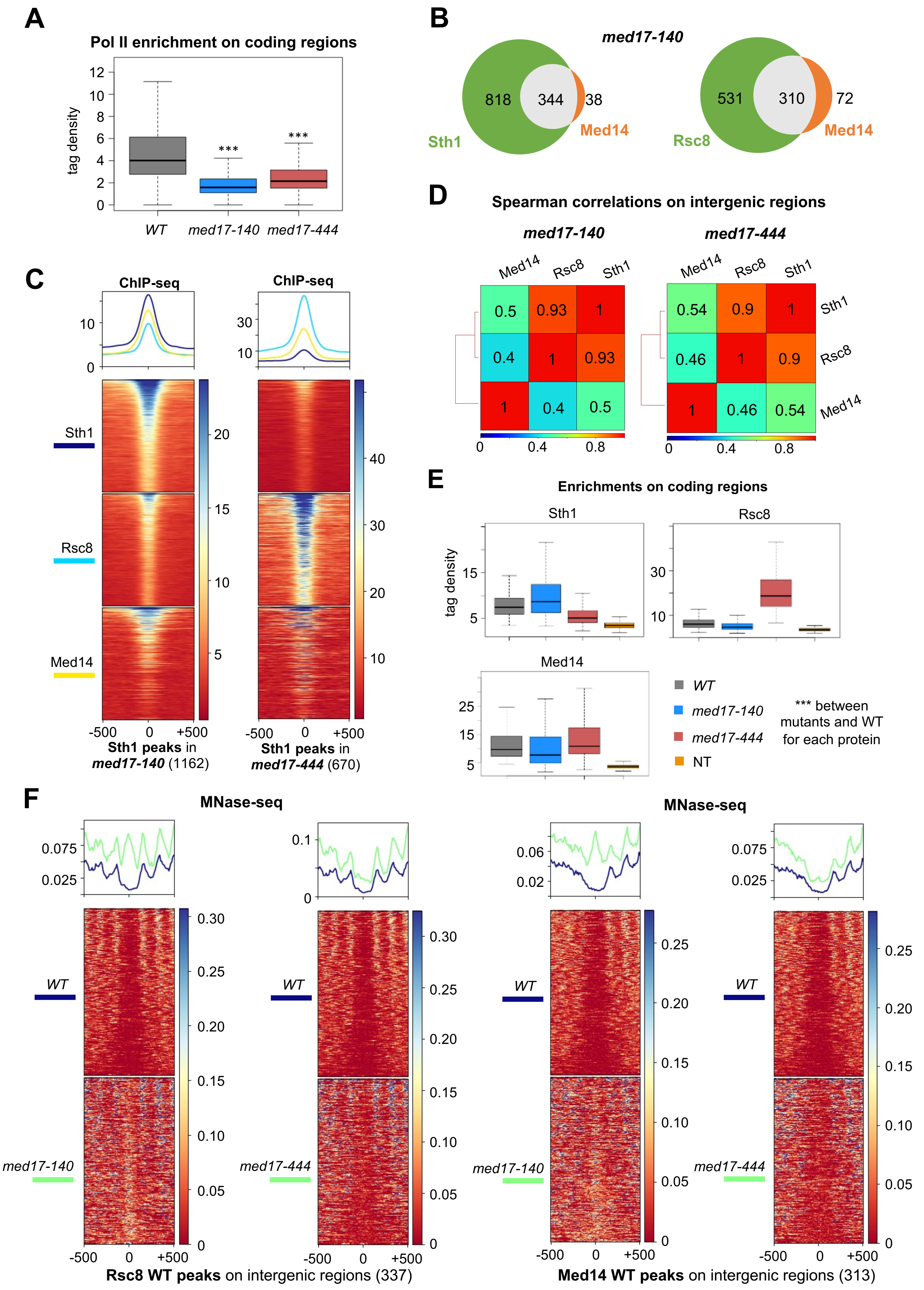

### Supplementary Figure S6

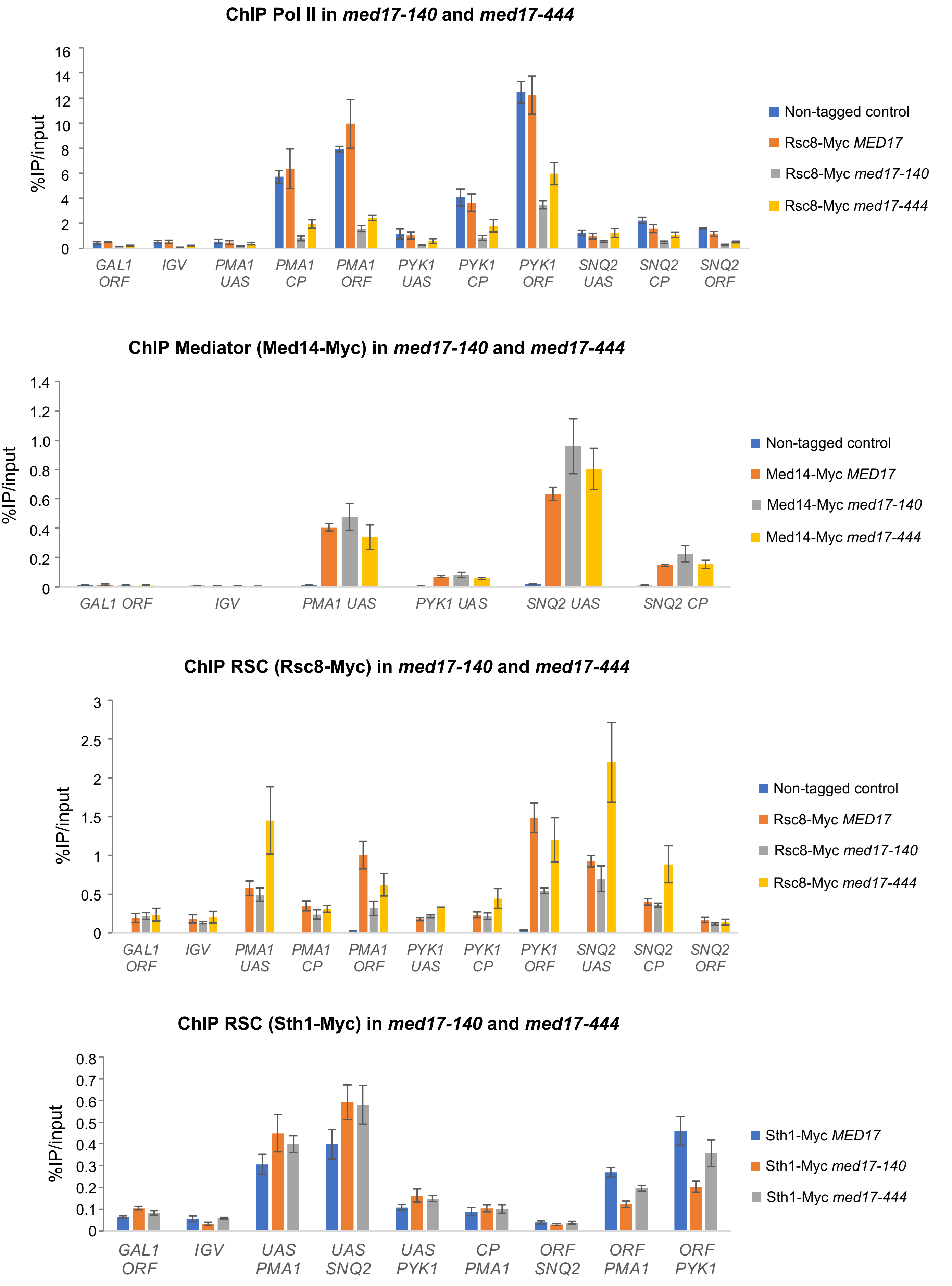

#### Supplementary Figure S7

A

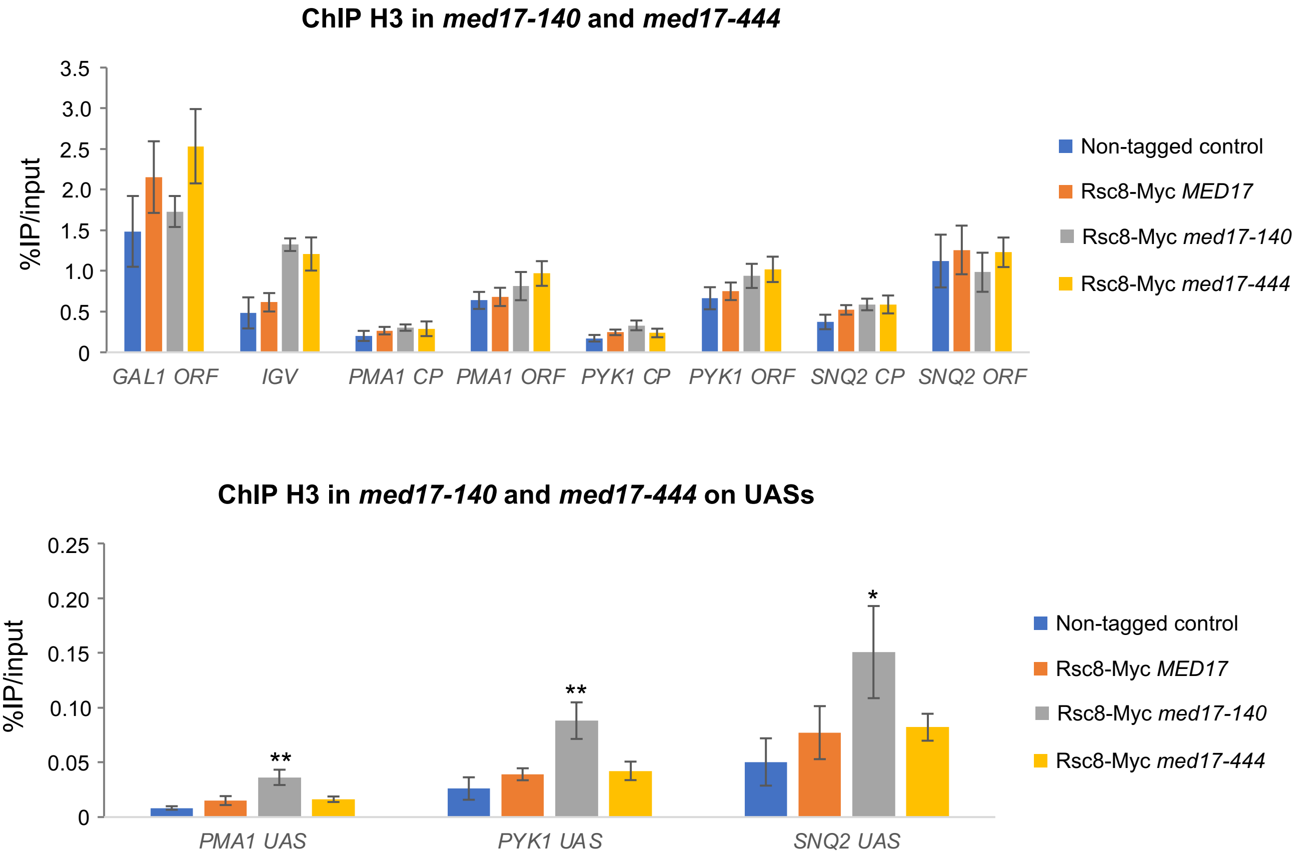


B

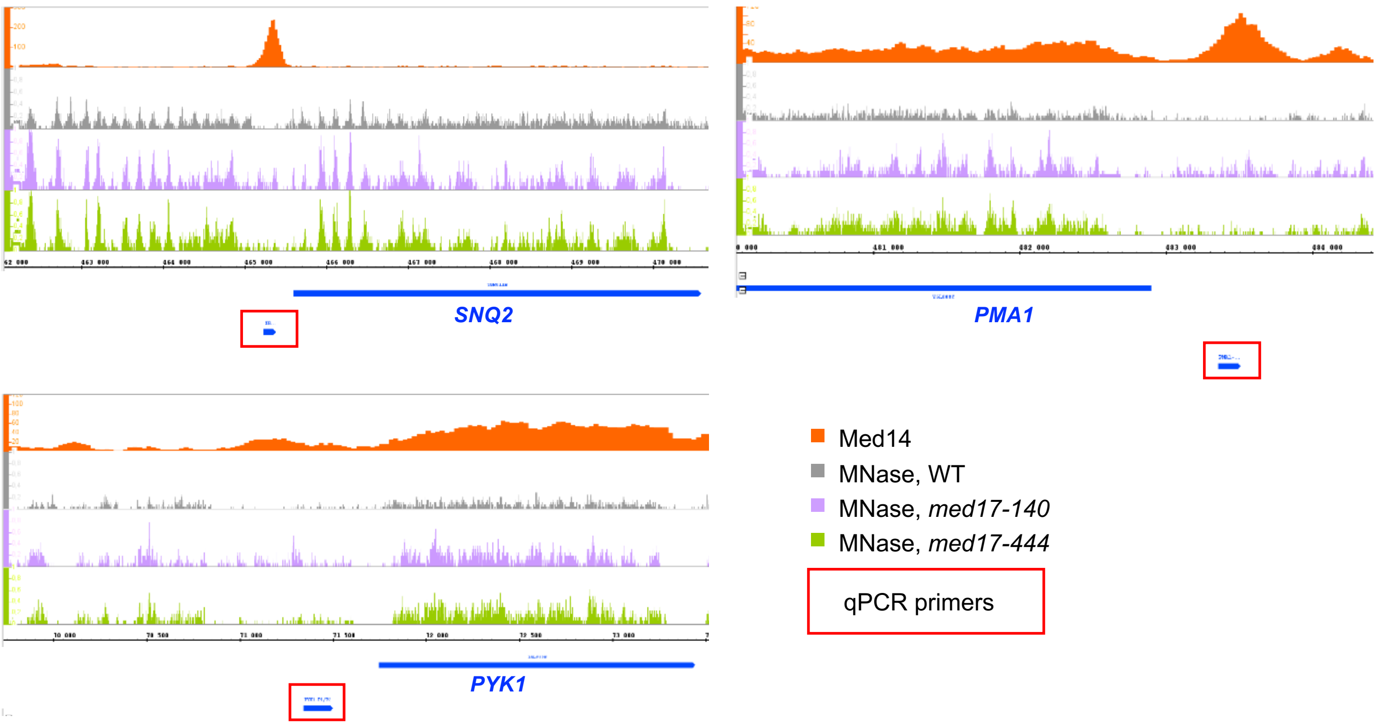

### Supplementary Figure S8

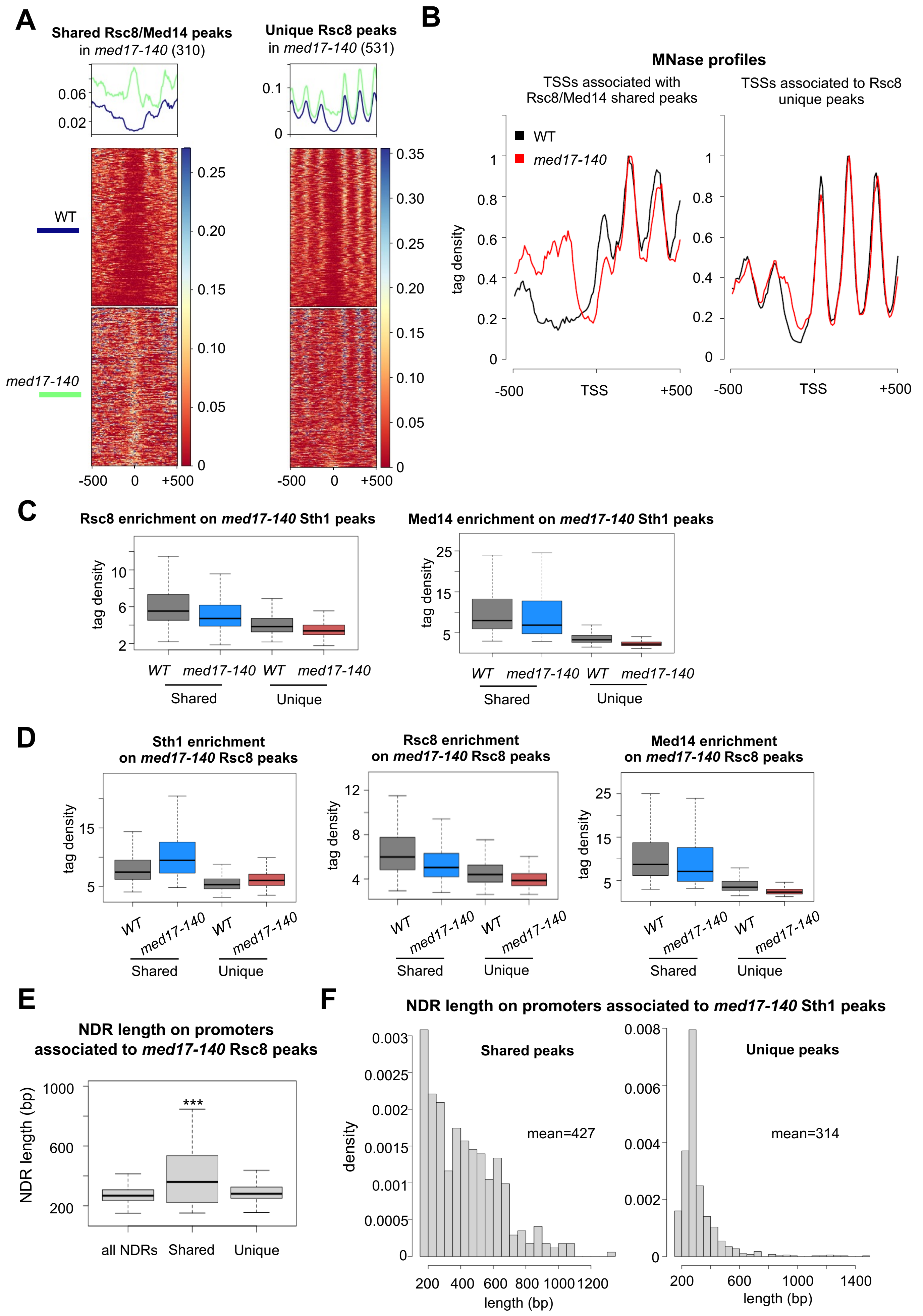

### Supplementary Figure S9

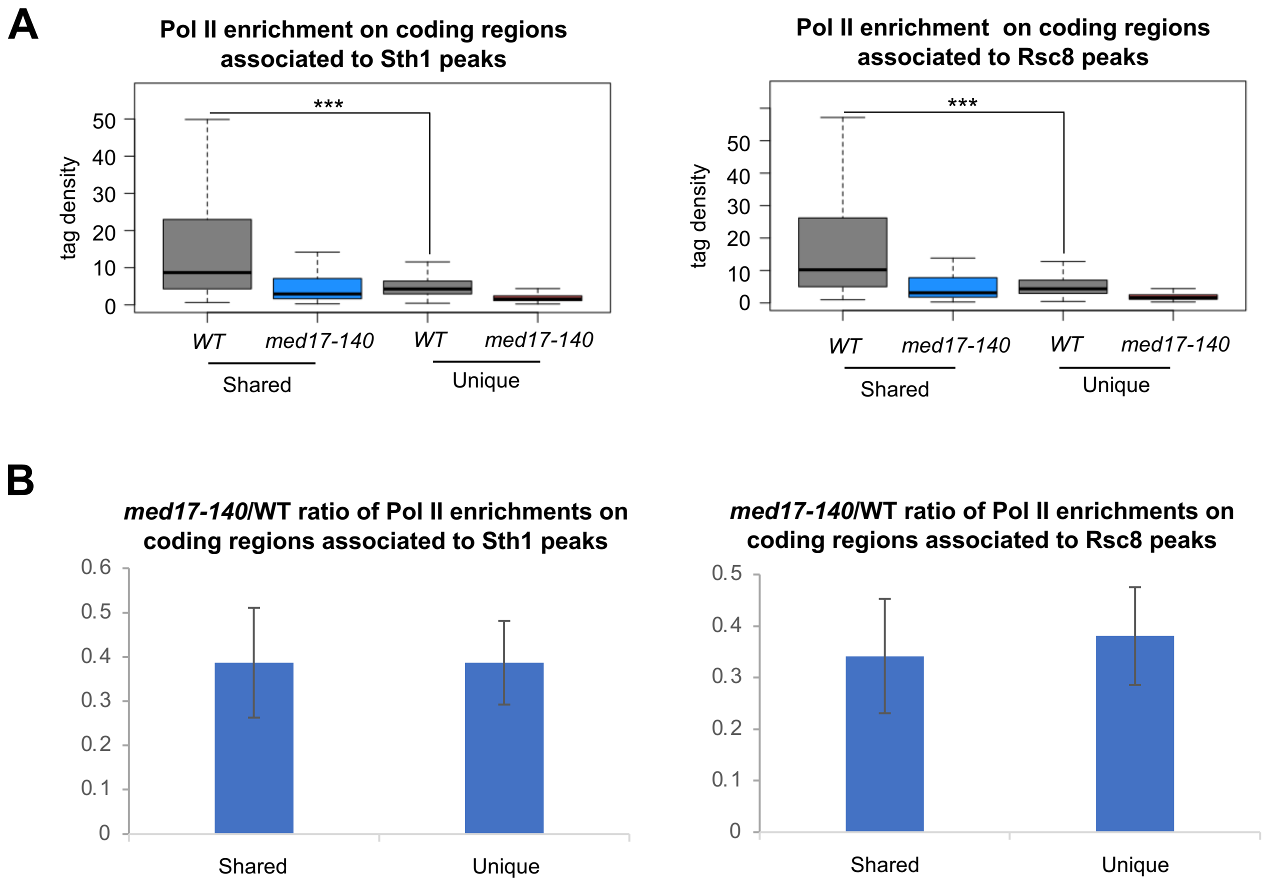
